# Generation of a biliary tract cancer cell line atlas reveals molecular subtypes and therapeutic targets

**DOI:** 10.1101/2024.07.04.601970

**Authors:** Vindhya Vijay, Negin Karisani, Lei Shi, Yu-Han Hung, Phuong Vu, Prabhat Kattel, Lauren Kenney, Joshua Merritt, Ramzi Adil, Qibiao Wu, Yuanli Zhen, Robert Morris, Johannes Kreuzer, Meena Kathiresan, Xcanda Ixchel Herrera Lopez, Haley Ellis, Ilaria Gritti, Lilian Lecorgne, Ines Farag, Alexandra Popa, William Shen, Hiroyuki Kato, Qin Xu, Eranga R. Balasooriya, Meng-Ju Wu, Saireudee Chaturantabut, Robin K. Kelley, James M. Cleary, Michael S. Lawrence, David Root, Cyril H. Benes, Vikram Deshpande, Dejan Juric, William R. Sellers, Cristina R. Ferrone, Wilhelm Haas, Francisca Vazquez, Gad Getz, Nabeel Bardeesy

## Abstract

Biliary tract cancers (BTCs) are a group of deadly malignancies encompassing intrahepatic and extrahepatic cholangiocarcinoma, gallbladder carcinoma, and ampullary carcinoma. Here, we present the integrative analysis of 63 BTC cell lines via multi-omics clustering and genome-scale CRISPR screens, providing a platform to illuminate BTC biology and inform therapeutic development. We identify dependencies broadly enriched in BTC compared to other cancers as well as dependencies selective to the anatomic subtypes. Notably, cholangiocarcinoma cell lines are stratified into distinct lineage subtypes based on biliary or dual biliary/hepatocyte marker signatures, associated with dependency on specific lineage survival factors. Transcriptional analysis of patient specimens demonstrates the prognostic significance of these lineage subtypes. Additionally, we delineate strategies to enhance targeted therapies or to overcome resistance in cell lines with key driver gene mutations. Furthermore, clustering based on dependencies and proteomics data elucidates unexpected functional relationships, including a BTC subgroup with partial squamous differentiation. Thus, this cell line atlas reveals potential therapeutic targets in molecularly defined BTCs, unveils biologically distinct disease subtypes, and offers a vital resource for BTC research.

## Introduction

Biliary tract cancers (BTCs) constitute a group of malignancies of the biliary tree epithelium anatomically categorized as intrahepatic cholangiocarcinoma (ICC), extrahepatic cholangiocarcinoma (ECC), gallbladder cancer (GBC), and ampullary/periampullary carcinoma (AC)^1^. Although relatively rare in North America and Europe, accounting for only 3% of all gastrointestinal malignancies, rates have been rising for decades (due primarily to ICC), and the incidence is much higher world-wide^2,3^. Moreover, BTC imposes a disproportionate public health burden, marked by an alarmingly low 5-year survival rate of <10% for advanced disease. While the discrete BTC subtypes are often grouped together clinically, with most patients with advanced disease receiving the same first-line chemotherapy, they exhibit extensive biological differences^4–6^, emphasizing the importance of better classification to guide treatment advances.

The marked molecular heterogeneity within and between the anatomical subtypes of BTC adds further complexity. In ICC, common genomic alterations involve the *IDH1*, *FGFR2*, *KRAS*, and *BRAF* oncogenes and the *TP53*, *BAP1*, *PBRM1*, and *ARID1* tumors suppressors^7–9^. In ECC, AC, and GBC, *TP53* and KRAS mutations predominate, and mutations in *SMAD4*, *ELF3* and WNT pathway genes (*APC*, *CTNNB1*, *AXIN1*) are enriched, although frequencies vary considerably depending on geography and etiology^5,7,10–13^. The genomics of BTC has guided precision medicine, with targeted therapies against *IDH1*, *FGFR2*, *BRAF*, and *ERBB2* alterations providing clinical benefit^4,14^. Despite these advances, it is crucial to enhance the modest increases in survival conferred by these therapies and to develop treatments against the larger proportion of patients lacking currently targetable alterations. These considerations underscore the imperative of establishing a functionally relevant molecular taxonomy for BTC, relating genomics and biological properties to each other and to clinical features. Such a system should enhance comprehension of similarities and differences among these tumors, identify essential pathways, and inform the design of treatments tailored to specific subsets of this disease.

Cell line models provide tractable systems for elucidating the oncogenic circuitry and spectrum of dependencies in various cancer types via large-scale functional genomics^15^. Given the diverse genomic and tumor biological features of BTCs, there is a pressing need to create and characterize a broad set of BTC cell line models that retain molecular faithfulness to the different subsets of patient tumors. In this study, we document the development of a comprehensive BTC cell line atlas. The primary objectives of this atlas are three-fold: first, to forge a collection of novel patient-derived BTC cell line models representing the diversity of genotypes and anatomical subtypes of the disease; second, to systematically chart the molecular features and vulnerabilities of BTC models; and third, to relate observations to clinical samples. Our study reveals novel dependencies associated with specific genotypes and cell lineage signatures, shedding light on previously unrecognized functional relationships, and introducing new, biologically significant subtypes of BTC. We envision this atlas and cell line collection as a primary resource for the BTC research community, fostering ongoing exploration and discovery.

## Results

### Establishment of a biliary tract cancer cell line collection representing the diverse features of the clinical disease

We compiled a collection of 63 BTC cell lines, including 30 new patient-derived cell line models generated using specimens obtained from biopsies, surgical resections, and rapid autopsies.

The collection represented the distinct anatomical subtypes of BTC, including ICC (n=40), GBC (n=10), ECC (n=8), and AC (n=5). Whole exome sequencing, RNA sequencing, and quantitative proteomics was performed on 56/63 cell lines, with the rest subjected to subsets of these assays (**Figure 1a** and **b, Table S1**). The library had broad representation of the heterogenous genomic profile observed in patient samples (**Figure 1c**). Consistent with patient data*, FGFR2* fusions and *IDH1*/*IDH2* mutations were restricted to ICC models and other trends included *KRAS* mutations in ECC and AC, *SMAD4* mutations in ECC, and *CTNNB1* mutations in AC. In addition, the arm-level copy number alterations (CNAs) observed in the cell lines largely aligned with the CNAs in patient samples, with significant overlap in chromosome 1p, 3p, 4q, 9p, and 14q losses and chromosome 11q gains (**Supplementary Figure 1a**).

**Figure 1.**
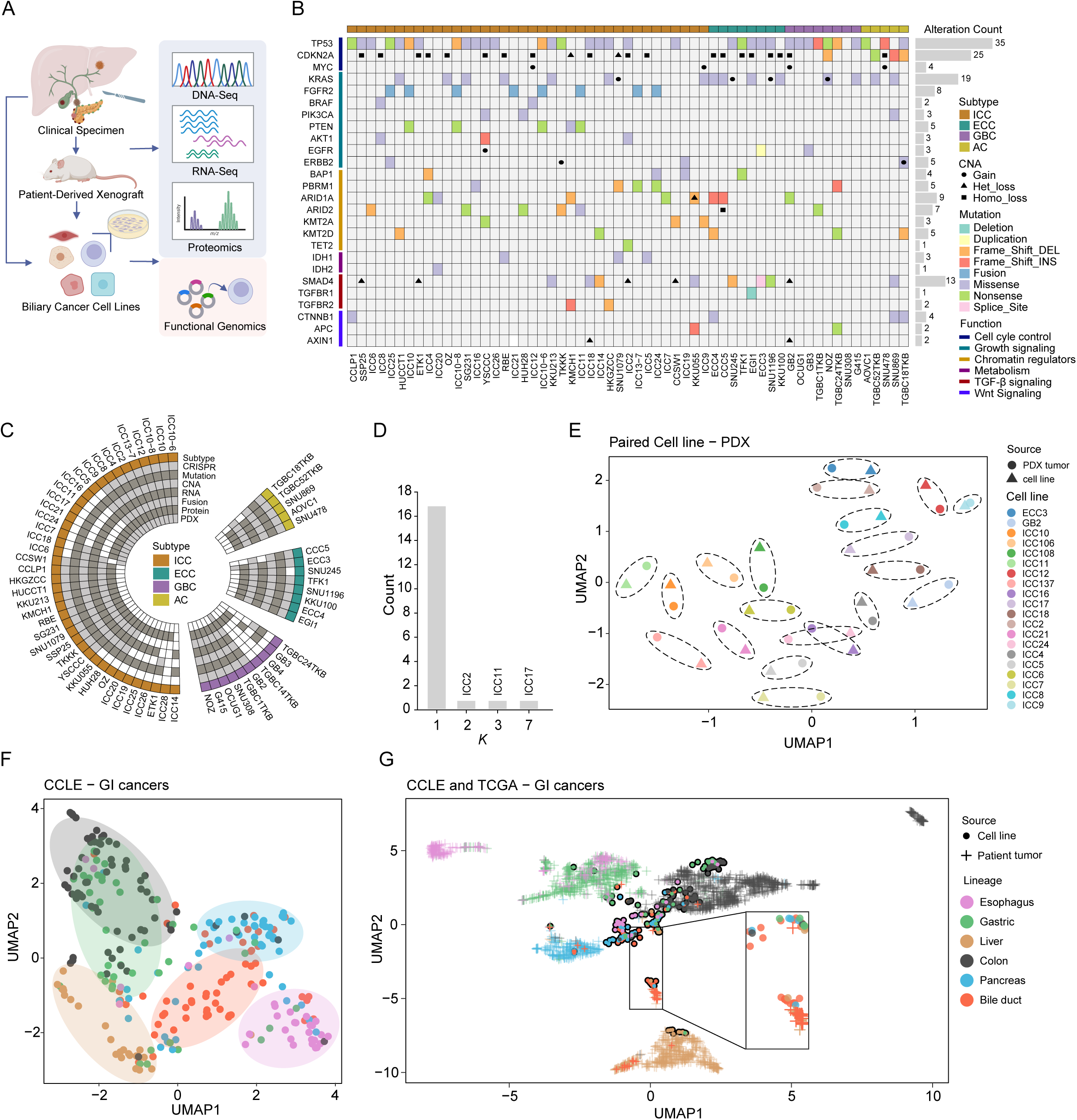
Molecular characterization of BTC cell line models. (A) Illustration of cell line model development and analysis approach. (B) Circular representation of BTC models included in the study (n=63), including ICC (n=40), ECC (n=8), GBC (n=10), and AC (n=5). Datasets available for each model are highlighted (grey). (C) Genomic characterization of BTC cell lines, including ICC (n=38), ECC (n=8), GBC (n=8), and AC (n=5). (D) Bar plot showing dispersion of K nearest cell line values for each cell line-PDX pair (n=20). (E) Two-dimensional UMAP representation of RNA sequencing data of cell lines (dots) and matched PDX tissues (triangles) (n=20). Cell line-PDX pairs are circled using a dotted line. (F) Two-dimensional UMAP representation of RNA sequencing data of BTC cell lines (n= 56) and cell lines derived from other DepMap gastrointestinal (GI) cancers (n=216), including Esophagus (n=32), Gastric (n=42), Liver (n=24), Colon (n=67), and Pancreas (n=51). Cell lines from each GI cancer type are colored as indicated. (G) Two-dimensional UMAP representation of RNA sequencing data of BTC cell lines (n=49) and other GI cancers (n=216), as well as corresponding primary tumors from TCGA data (n=1,974) representing BTC (n=45) and other GI cancers, including Esophagus (n=196), Gastric (n=450), Liver (n=423), Colon (n=677), and Pancreas (n=183).

A key question was whether these *in vitro* models retain the molecular circuitry of the tumor types from which they were derived. To address this, we first compared the transcriptomes of 20 of the cell lines with matched patient-derived xenograft (PDX) models (established directly from patient tumors without *in vitro* culture). By calculating the Euclidean distance between the PDXs and cell lines, we determined the K^th^ nearest neighbor for each model. 17/20 (85%) of the cell lines were the nearest neighbors with their respective PDXs, while 3/20 had slight deviations, where ICC17 was the most distant pair, whereas ICC11 and ICC2, although not being the 1^st^ nearest neighbors had similar distance and correlation as the nearest neighboring pairs (**Figure 1d, Supplementary Figure 1b**). This similarity is also evident in the two-dimensional projection using the Uniform Manifold Approximation and Projection (UMAP) analysis (**Figure 1e**). We next compared our library of BTC cell lines with 216 cell lines derived from other types of gastrointestinal (GI) cancer in the DepMap database^15^. The BTC cell lines showed a distinct clustering (in UMAP), projecting between hepatocellular carcinoma (HCC; Liver) and pancreatic cancer cell lines, consistent with the common origins of these lineages from the foregut endoderm (**Figure 1f**). Extension of this correlative analysis to TCGA patient data representing GI cancers showed that the majority of the BTC cell lines clustered tightly with BTC patient specimens, with the exceptions mainly grouping with pancreatic cancers and HCCs (**Figure 1g**). Thus, despite adaptation to 2D culture and passaging *in vitro*, most of the BTC cell lines broadly maintain transcriptional programs characteristic of the primary tumors.

### Mapping genotype-specific genetic vulnerabilities in BTC

Next, we characterized the genetic vulnerabilities of 39 of the BTC cell lines via genome-wide CRISPR loss-of-function screens that we incorporated into the Broad Cancer DepMap (>1000 cell lines representing cancers from > 30 different tissues). Here, we considered any gene with a Chronos gene effect score of <-0.5 as a dependency (Methods). The multi-dimensional datasets provided several organizing principles to uncover and understand differences and similarities between BTCs, relating dependencies variously to genetic, anatomic, transcriptional, and proteomic features.

As a first level of analysis, we studied the dependencies specifically associated with the recurrent genomic alterations found in BTC. Multiple models within our library harboring oncogenic mutations in established therapeutic targets^4,16^ were screened. Four cell lines were derived from patients with *FGFR2*-fusion+ ICC who progressed on FGFR inhibitor therapy (ICC13-7, ICC10, ICC10-6 and ICC10-8). These cell lines exhibited an intermediate level of genetic dependency on *FGFR2* (with the exception of ICC10-8, which showed loss of expression of the FGFR2 fusion), matching our previous studies using pharmacological FGFR inhibition (**Figure 2a**, and see^17^). BTC lines with BRAF-V600E mutations (ICC8, ICC12) were among the most highly *BRAF*-dependent models within the DepMap collection (**Figure 2b**), whereas other variants of *BRAF* (L485W and I775F) did not confer dependency (**Supplementary Figure 2a**). *ERBB2* was essential in BTC cell lines with amplification (TKKK cells) or activating mutations (S310F and R678Q in TGBC18TKB cells) (**Supplementary Figure 2b)**. By contrast, mutant *IDH1* (R132C, SNU1079 cells; R132S, RBE cells) did not confer dependency (**Figure 2c**), in line with evidence that this oncogene does not function as a conventional driver of proliferation or survival in solid tumor cells *in vitro*^18,19^. Additionally, BTC cell lines with activating mutations in *KRAS* (at codons 12, 13, 61, N= 14 models), *PIK3CA* (E545A, E545K, N=2), and *CTNNB1* (G34V, S33P, T41A, S45P, N=4) displayed strong dependencies on the respective gene (**Figure 2d-f**). These findings reveal prominent oncogene addiction for important BTC genotypes and suggest that an increasing proportion of patients with BTC may respond to current or emerging targeted therapies.

**Figure 2.**
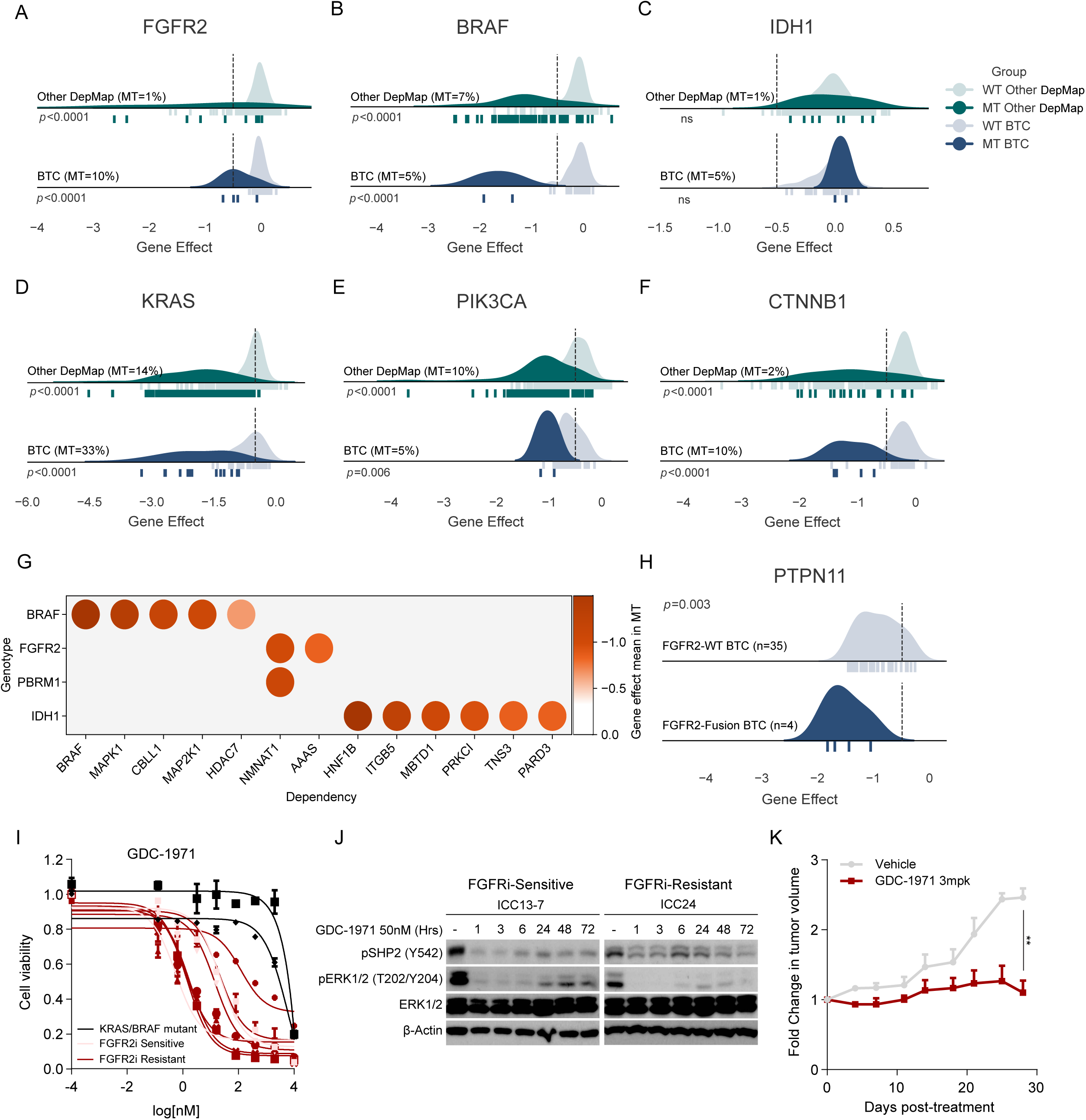
Identification of genotype-specific vulnerabilities. (A-F) Density plots of gene effect scores (Chronos scores) in BTC (n=39) and non-BTC cell lines (n=1061) harboring genetic alterations as indicated (rate in %). Dotted lines show the -0.5 dependency threshold. Unpaired t-test comparing dependency of wild-type (WT) vs. mutant (MT) at each level. (G) Dot plot showing key significant differential gene dependencies associated with each genetic subset. Unpaired t-test; adjusted p < 0.1. (H) Density plot showing distribution of PTPN11 gene effect in FGFR2-fusion+ (n=4) versus wildtype BTC cell lines (n=35). Dotted lines indicate the -0.5 dependency threshold. Unpaired t- test. (I) Drug-response curves for GDC-1971 in BTC cell lines. Data are shown as mean ± SEM. Black= KRAS/BRAF mutant ICC, Pink= FGFR2i sensitive ICC and Red= FGFR2i resistant ICC cell lines. (J) Immunoblot analysis of the indicated signaling proteins in ICC13-7 (FGFRi-sensitive) and ICC24 (FGFR2i-resistant) cells upon treatment with 50nmol/L GDC-1971 for indicated time. (K) Growth curve showing fold change in tumor volume of ICC11 cells implanted subcutaneously. Fold change was calculated by normalizing to the initial tumor volume. Data is shown as mean ± SEM. Unpaired t-test

### Synthetic lethal relationships in genetic subsets of BTC

Next, we sought to identify other genetic vulnerabilities associated with oncogene activation or tumor suppressor loss. No strong differentially enriched vulnerabilities were associated with genomic alterations of *TP53*, *ARID1A or SMAD4*, suggesting extensive heterogeneity within these groups. Similarly, cell lines with mutations in *KRAS* and *CTNNB1*, were only differentially dependent on *KRAS* and *CTNNB1*, respectively, but not on any other gene (**Supplementary Table S2**). By contrast, cell lines with *BRAF* V600E mutations were enriched for dependencies on the downstream MEK/ERK pathway genes, MAP2K1 and MAPK1, E3 ubiquitin ligase CBLL1, and the type IIA histone deacetylase family member, HDAC7, among others (**Figure 2g, Supplementary Figure 2c**). Cell lines with FGFR2 fusions or *PBRM1* loss were highly dependent on the NAD biosynthetic enzyme, NMNAT1 (**Figure 2g, Supplementary Figure 2d, e)**. *IDH1* mutant cell lines showed a number of notable dependencies. These included MBTD1, a methylated histone H4K20 reader protein, potentially reflecting changes in epigenetic regulation emerging from the aberrant increase in histone methylation caused by mutant IDH1^19,20^.The HNF1B transcription factor and a set of cell polarity proteins (ITGB5, PRKCI, TNS3, and PARD3) were also distinct vulnerabilities in these cell lines (**Figure 2g, Supplementary Figure 2f**). Importantly, each of these dependencies was strongly enriched across all *IDH1* mutant cell lines from epithelial cancers in the DepMap (**Supplementary Figure 2g**). Collectively, these data point to new features of the cellular circuitry driving the growth of important genetic subsets of BTC, including some pathways that may be targetable therapeutically (e.g., PRKCI).

PRMT5 and PTPN11 are additional emerging precision medicine targets. Inactivation of the PRTM5 arginine methyltransferase has been identified as synthetic lethal with deletion of methylthioadenosine phosphorylase (MTAP), which is frequently co-deleted with the neighboring CDKN2A tumor suppressor^21,22^. CDKN2A/MTAP-deleted BTC cell lines showed prominent PRMT5 dependency (**Supplementary Figure 2i**), supporting the further clinical exploration of PRMT5 inhibitors in MTAP-deleted BTCs. PTPN11 encodes SHP2 protein phosphatase that links receptor tyrosine kinase (RTK) signaling and RAS activation. SHP2 inhibitors are in clinical trials against tumors with RTK/mitogen-activated protein kinase (MAPK) pathway alterations^23^. This strategy may be particularly relevant in FGFR2-fusion+ ICCs, which exhibit enriched SHP2 activity^24^ and show FGFR2-dependent SHP2 regulation in associated models^17^. Moreover, SHP2 signaling is sustained in models from patients who developed FGFR inhibitor resistance^17,25^. FGFR2-fusion+ cell lines were highly dependent on PTPN11 (**Figure 2h and Supplementary Figure 2j**). Extending these findings, the allosteric SHP2 inhibitor, GDC- 1971, demonstrated prominent activity (IC50 <120 nM) across 8 FGFR2-fusion+ ICC lines that are either sensitive/partially sensitive or resistant to FGFR inhibitor treatment; by contrast G415 (KRAS-G13D mutant) and ICC12 cells (BRAF-V600E mutant) were resistant (IC50 > 1000 nM) (**Figure 2i**). Western blots confirmed the suppression of downstream ERK signaling (**Fig. 2j** and **Supplementary Figure 2k, l**). Finally, GDC-1971 demonstrated efficacy *in vivo* against the FGFR TKI-resistant ICC11 xenograft model^17^ (**Figure 2k**). Together, these data support the testing of SHP2 inhibitors in FGFR2-fusion+ ICC, particularly in the setting of FGFR inhibitor resistance.

### EGFR is a selective vulnerability in BTC, associated with chromosome 3p loss

The generation of a dependency map (DepMap) enables comparison of the circuits broadly required in BTC compared to cell lines from other cancer types. It also permits the unbiased identification of functionally related groups of BTC cell lines that extend beyond classification based solely on single gene mutations. Accordingly, we sought to capture shared features and distinctive patterns present across our BTC cell lines by first computing the top 20 most preferential dependencies in each model (encompassed 696 unique genes, **Table S3**). We next asked whether specific pathway requirements distinguish BTC cell lines and cell lines from other solid tumor types. Comparison of the profile of BTC dependencies with those in pancreatic adenocarcinoma (PDAC, n=61) (n=1045 unique genes) and HCC, n=21 (n=388 unique genes) revealed limited overlap across cancer types (**Supplementary Figure 3a, b**). Pathway Enrichment Analysis demonstrated some functional convergence (*e.g*., adherens junction pathway), as well as selective requirements of individual tumor types, with BTC showing specific enrichment of the actin cytoskeletal dynamics, central carbon metabolism, and EGFR signaling pathways (**Supplementary Figure 3b**). Collectively this analysis points to extensive differences in oncogenic circuitry between gastrointestinal tumor types.

This prompted us to define specific dependencies enriched in BTC relative to the complete DepMap collection. The most significant BTC-associated dependencies were EGFR and the RAC/RHO guanine nucleotide exchange factors, DOCK5, and ARHGEF7, which integrate RTK signaling and cytoskeletal changes. (**Figure 3a**). The two BTC cell lines in our library with EGFR kinase domain mutations (TGBC1TKB cells, EGFR-L883F and ECC3 cells, -insS768_D770dup) and one with an amplification (YSCCC cells) each showed EGFR dependence (gene effect score <-0.5) (**Figure 3b**). However, several of the most EGFR-dependent BTC cell lines lacked genomic alterations of EGFR. Overall, 64% (25/39) of BTC lines were dependent on EGFR (score <-0.5), versus 23% of other cell lines (**Supplementary Figure 3c**).

**Figure 3.**
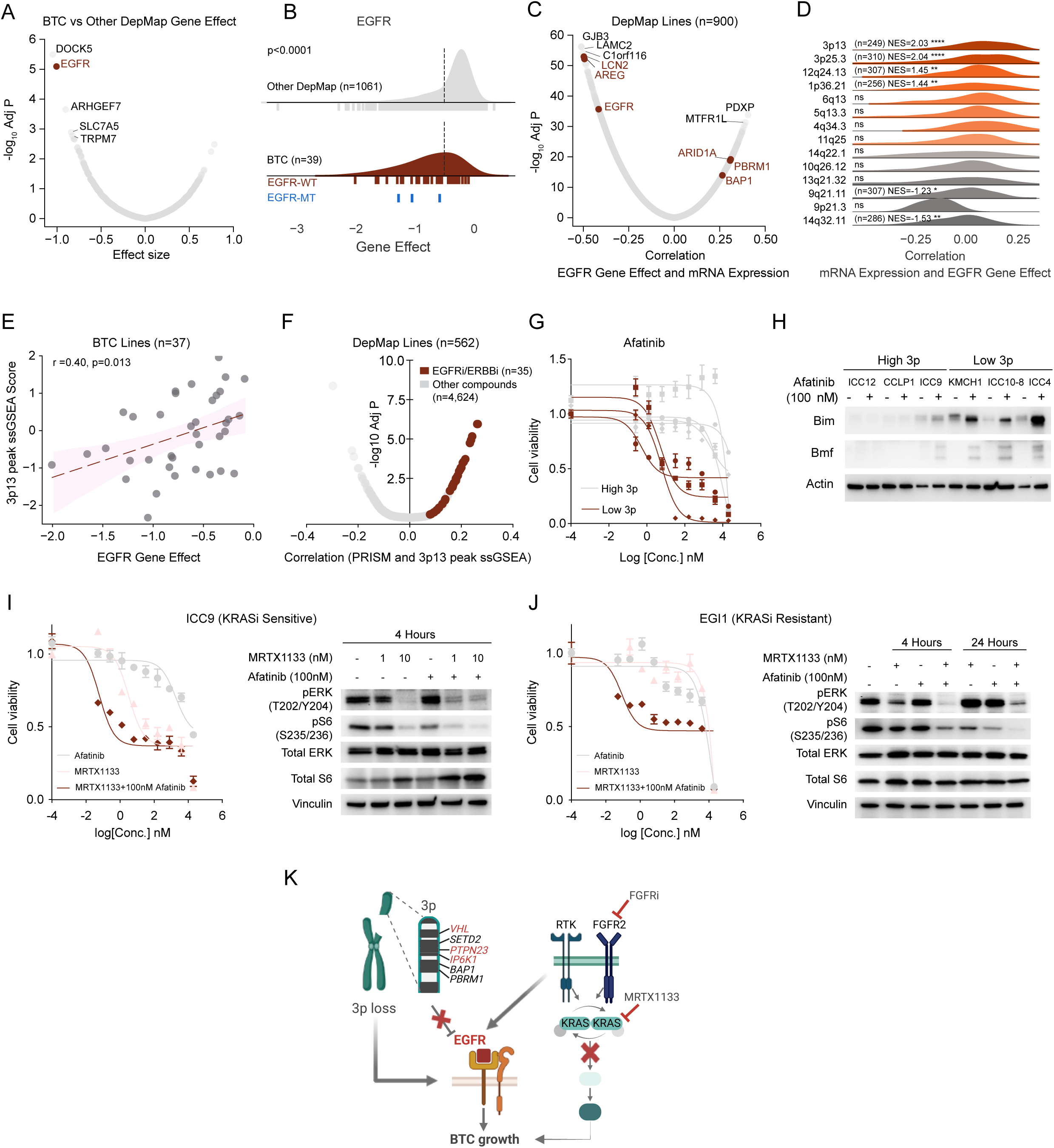
EGFR dependency is a selective vulnerability in BTC. (A) Volcano plot showing differential dependencies comparing BTC cell lines (n=39) and non-BTC (n=1020) cell lines. Unpaired t-test. (B) Density plot showing distribution of EGFR gene effect in BTC (brown) and non-BTC (grey) cell lines. Dotted lines indicate the -0.5 dependency threshold. Unpaired t-test. (C) Volcano plot showing correlation of gene expression with EGFR dependency across DepMap. Pearson correlation test. (D) Density plots showing distribution of pearson coefficients corresponding to correlation between EGFR gene effect and mRNA expression. Genes were grouped by arm level GISTIC peaks. FGSEA, NES, normalized enrichment score; ns, not significant; adjusted p < 0.1 (see methods). (E) Scatter plot showing correlation of EGFR gene effect and 3p13 peaks ssGSEA score of all BTC cell lines (n=39). Pearson correlation r; Wald test p with t-distribution. (F) Volcano plot showing correlation of 3p13 peaks ssGSEA score and response to inhibitors (n=568) in the DepMap PRISM repurposing screen across DepMap (n=562). EGFR/ERBB2 inhibitors (n=35) are highlighted in brown. Pearson correlation test. (G) Dose-response curves for afatinib in 3p13 low (red) cell lines (n=3) and 3p13 high (grey) (n=4). Data are shown as mean ± SEM. (H) Immunoblot analysis of the indicated apoptosis signaling proteins in high and low 3p13 scoring cell lines treated with afatinib at 100 nmol/L or DMSO (vehicle) for 4 hours. (I-J) (Left) Dose-response curves for EGFR inhibitor Afatinib (grey), KRAS^(G12D)^ inhibitor MRTX1133 (pink) or combination of 100nM Afatinib and MRTX1133 (red) in KRASi sensitive cell line ICC9 (I) or resistant cell line EGI1 (J). (Right) Immunoblot analysis of the indicated signaling proteins in ICC9 (I) or EGI1 (J) when treated as indicated for 4 or 24 hours

We next examined transcriptomic predictors of EGFR dependency and found high correlations with the expression levels of the EGFR trafficking factor, LCN2 within BTC and across the DepMap (**Figure 3c** and **Supplementary Figure 3d**), and with MAPK/JNK regulator, MAP3K8, specifically in BTC (**Supplementary Figure 3e, f**). EGFR dependency also correlated with low expression of the PBRM1 and BAP1 tumor suppressors (**Figure 3c**), which are located on chromosome 3p and are often deleted or mutated in BTC^26^. Accordingly, evaluation of the different CNA regions commonly altered in BTC (from TCGA) revealed that loss of 3p13 peak genes correlated with EGFR dependency (**Figure 3d, e**; see Methods) and EGFR mRNA expression (**Supplementary Figure 3g**). Notably, these associations with 3p loss were also evident across the DepMap (**Supplementary Figure 3h, i)**. Furthermore, we observed a striking association between EGFR/ERBB inhibitor sensitivity and 3p loss in the PRISM screen^27^, which tested 4,686 anti-cancer compounds against 562 cancer cell lines (**Figure 3f**, see **Methods**).

Importantly, treatment of a series of EGFR wild type BTC cell lines with the selective EGFR/ERBB2 TKI, afatinib, showed that those with 3p loss (ICC4, ICC10-8 and KMCH1; below average 3p13 peak ssGSEA score) were highly sensitive compared to those with high 3p scores (CCLP1, ICC12, ICC9 and SNU1079; above average 3p13 peak ssGSEA score) (mean IC50 ∼4 nM versus mean IC50 >1000 nM) (**Figure 3g**). Western blot analysis demonstrated induction of the proapoptotic BIM and BMF proteins^17^ in the 3p deleted models (**Figure 3h**), associated with suppression of downstream MEK/ERK and mTOR/S6K signaling (**Supplementary Figure 3j**). Collectively, the data highlight the enriched dependency of BTC on EGFR and provide rationale for targeting EGFR in defined subsets of BTC that extend beyond those with genomic alterations in the pathway. These features (3p deletion, high expression of EGFR, LCN2 and MAP3K8) typically coincided, suggesting a general program driving EGFR dependency. Notably, 3p spans the VHL, PTPN23 and IP6K1 genes, which have been linked to the negative regulation of EGFR signaling in other contexts^28–30^.

### EGFR inhibition sensitizes BTCs to concurrent KRAS/FGFR2 inhibition

These findings motivated us to assess whether BTC cell lines rely on the EGFR pathway in additional biological contexts. In this regard, previous work showed that the efficacy of FGFR inhibitors against FGFR2-fusion+ ICC cell lines and PDX models is potentiated by combined treatment with an EGFR/ERBB TKI, which overcomes feedback reactivation of MEK/ERK signaling^17^. To investigate whether this strategy can be extended to other targeted therapies, we tested the combined impact of the selective KRAS^G12D^ inhibitor, MRTX1133, and afatinib against KRAS-G12D mutant BTC cell lines. In both ICC9 (sensitive to MRTX1133 [IC_50_=3 nM] and resistant to afatinib [IC_50_ > 1000 nM]) and EGI1 cells (resistant to both [IC50 > 10 µM]), combination treatment with 100 nM afatinib was strongly cooperative (MRTX1133 IC_50_ = 0.06 nM and IC_50_ = 0.08 nM, respectively) (**Figure 3i, j**). These results support the use of EGFR/ERBB inhibitors to potentiate mutant KRAS targeting in BTC, further reinforcing the importance of EGFR in this malignancy (**Figure 3k**).

### Identification of selective dependencies in anatomical subtypes of BTC

The different types of biliary epithelia originate from multipotent hepato-pancreato-biliary progenitors in the foregut endoderm but have distinct developmental trajectories^31,32^. The intrahepatic bile ducts emerge from bipotential hepatoblast progenitors that also give rise to the hepatocytes. The other parts of the biliary tree arise from budding of an adjacent region of the gut tube. Accordingly, we sought to explore potential selective dependencies between the different anatomical subtypes of BTC. We included HCC in these studies given the relatedness of hepatobiliary tissues. Differential dependency analysis between each subtype and the other cell lines identified up to 6 significantly selective dependencies, with strong selective dependencies for GBC, ECC, and ACC cell lines (**Figure 4a, Supplementary Figure 4a and Table S4**).

**Figure 4.**
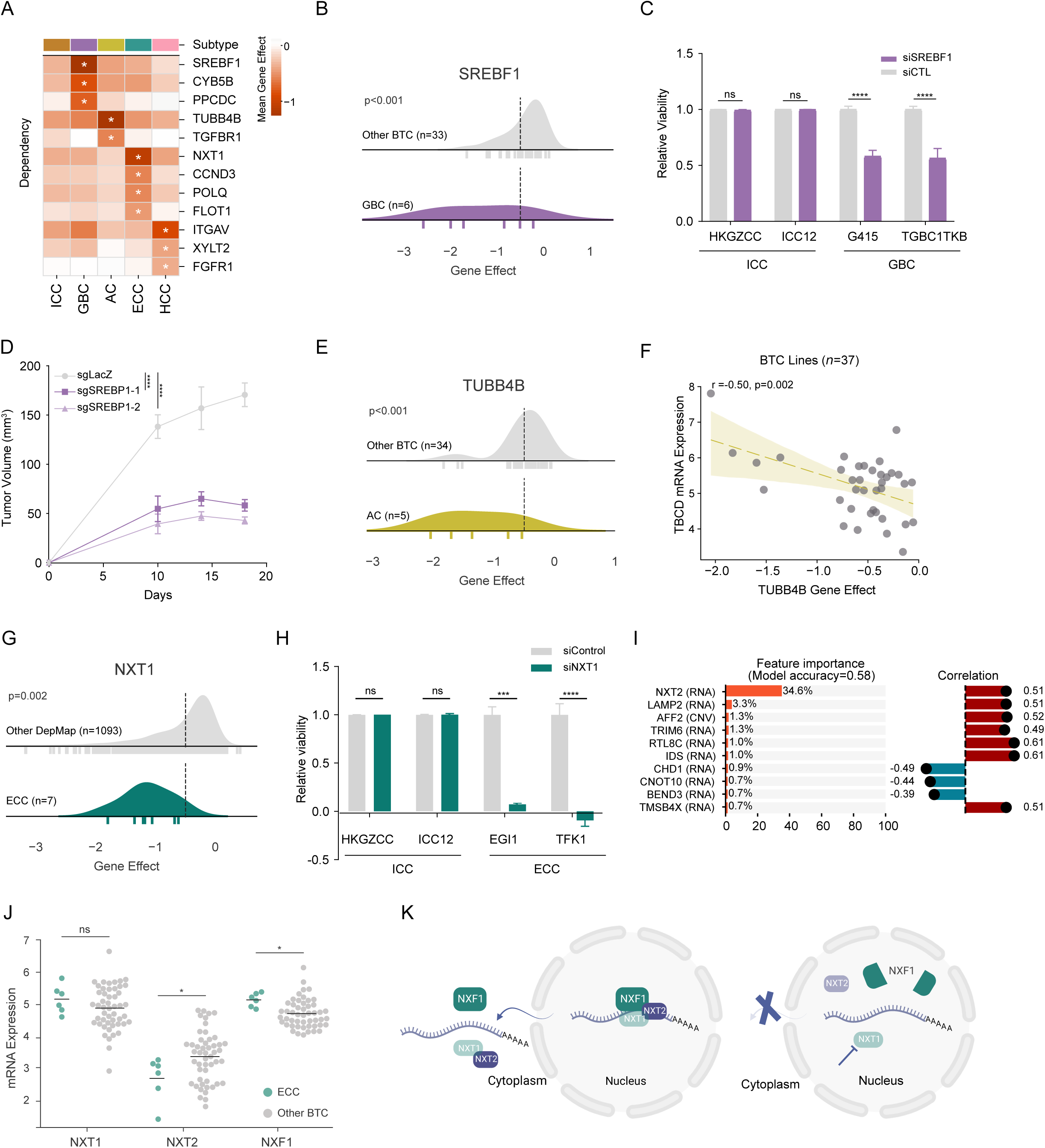
Selective dependencies associated with ECC and GBC. (A) Heatmap indicating selective dependencies associated with each anatomical subset of BTC. Unpaired t-test; *adjusted p <0.1. (B) Density plot showing SREBF1 dependency in GBC (n=6) or other BTC (n=33) cell lines. Dotted lines indicate the -0.5 dependency threshold. Unpaired t-test. (C) Bar plot showing relative viability of GBC or ICC cell lines after knockdown of SREBP1. Bar plots show mean ± SEM. Unpaired t-test. ns, not significant; ****p<0.001 (D) Serial measurements of tumor volume in mice sub-cutaneously injected with SNU308 cell line infected with sgLacZ (control) or sgSREBP1. (E) Density plot showing TUBB4B dependency in AC (n=5) or other BTC (n=34) cell lines. Dotted lines indicate the -0.5 dependency threshold. Unpaired t-test. (F) Scatter plot showing correlation between TUBB4B gene effect and TBCD mRNA expression in BTC cell lines (n=37). Pearson correlation r; Wald test p with t-distribution. (G) Density plot showing NXT1 dependency in ECC (n=8) and non ECC (n=31) BTC cell lines. Dotted lines indicate the -0.5 dependency threshold. Unpaired t-test. (H) Bar plots showing relative viability of ECC or ICC cell lines after NXT1 knockdown. Bar plots show mean ± SEM. Unpaired t-test. ns, not significant; ***p<0.001; ****p<0.0001. (I) Random Forest analysis showing the top 10 molecular features predictive of NXT1 dependency, along with their Pearson correlation with the NXT1 gene effect score among BTC cell lines. (J) Swarm plot showing NXT1, NXT2 and NXF1 mRNA levels in ECC or other BTC cell lines. Horizontal line segments indicate the mean. Wilcoxon rank-sum test; ns, not significant; *p<0.01 (K) Schematic showing relationship between NXT1, NXT2 and NXF1.

GBC cell lines exhibited strong selective dependency on PPCDC, required for coenzyme A (CoA) biosynthesis, CYB5B, a mitochondrial cytochrome reductase involved in fatty acid synthesis, and on SREBF1, the master transcriptional regulator of cholesterol metabolism. PPCDC stood out as a strongly selective dependency both relative to other BTC cell lines (4/5 of the most dependent BTC lines were GBC) and across the DepMap (**Supplementary Figure 4b**). Similarly, out of the 39 BTC lines screened, the top three SREBF1-dependent lines were GBCs, while the remaining three GBC cell lines showed intermediate dependency (**Figure 4b**). Moreover, GBC cell lines ranked first, third, and ninth in SREBF1 dependency score in the context of the entire DepMap (**Supplementary Figure 4c**). We confirmed that the GBC cell lines, TGBC1TK and SNU308, were highly sensitive to depletion of SREBF1 while both ICC cell lines tested as controls were unaffected (**Figure 4c**). Furthermore, SREBF1 knockout impaired subcutaneous tumor growth of SNU308 xenografts (**Figure 4d** and **Supplementary Figure 4d)**. SREBF1 dependency correlated with upregulation of the SREBF1 transcriptional target, MVK (**Supplementary Figure 4e**), providing a potential biomarker for this dependency. Since CoA carries activated acyl groups required for fatty acid catabolism and anabolism, the selective requirement of GBCs on PPCDC, CYB5B, and SREBF1 suggest that this tumor type might have an enhanced demand for lipid synthesis.

The most prominent differential dependency in AC was TUBB4B (**Figure 4e**), correlating with high expression of β-Tubulin polymerization cofactor D, TBCD (**Figure 4f**). Comparison of AC cells with the rest of the DepMap highlighted additional selective dependencies. This included riboflavin kinase (RFK) (**Supplementary Figure 4f**) and the large neutral amino acid transporter, SLC7A5, correlating with increased expression of its paralog, SLC7A8 (**Supplementary Figure 4g**). Overall, these enrichments in AC and GBCs indicate that there may be subtype-specific metabolic requirements among BTCs.

ECC cell lines showed strongly enriched dependency on the mRNA nuclear export factor, NXT1^33^ (**Figure 4g)**. Validating these results, siRNA-mediated knockdown of NXT1 blocked the growth of the ECC cell lines, EGI1 and TFK1, while not significantly affecting the ICC cell lines, HKGZCC and ICC12 (**Figure 4h** and **Supplementary Figure 4h**). NXT1 dependency correlated with low expression of the paralogous NXT2 gene (**Figure 4i** and **Supplementary Figure 4i**), as previously observed in neuroblastoma cells^34^. As in neuroblastoma, inactivation of NXT1 destabilized the pan-essential mRNA export factor, NXF1, selectively in NXT2-low ECC cell lines (**Supplementary Figure 4j**). Additionally, ECC cell lines showed significantly higher relative levels of NXF1 (**Figure 4j**), suggesting that they may have greater demand for the mRNA nuclear export machinery (**Figure 4k**). Thus, NXT1 may be a relevant target for the treatment of ECC. The clinical development of the XPO1 inhibitor, selinexor^35^, supports the therapeutic potential of targeting the nuclear transport machinery.

### Liver lineage features identify distinct CCA subtypes showing dependency on lineage survival factors

As an additional approach to identify novel, functionally significant relationships, we stratified the BTC cell lines based on their transcriptomic profiles. As above, we included HCC models in this analysis. Unsupervised clustering identified four RNA subgroups (**Figure 5a, Supplementary Figure 5a** and **Table S5**). HCC cell lines were primarily in clusters R1 and R2; ICC were primarily in R3 and R4; and ECC, GBC, and AC were mainly in R1 and R4. Random index analysis revealed that cluster R3 was enriched for genomic alterations in *FGFR2*, *IDH1*, *ARID1A*, *PBRM1* and *BAP1* and for wild type TP53 status, while R4 was enriched for alterations in *KRAS*, *BRAF, SMAD4*, CDKN2A, EGFR and ERBB2 (**Supplementary Figure 5b**). Pathway analysis demonstrated signatures for epithelial-mesenchymal transition in R1, fibrinolysis and cholesterol metabolism in R2, bile secretion and fatty acid biosynthesis in R3, and epithelial cell differentiation and vesicular transport in R4 (**Figure 5a**). Notably, we observed that R3 shared transcriptional features with both R2 (HCC-enriched) and R4, as determined by calculation of the correlation coefficient between clusters (**Supplementary Figure 5c**).

**Figure 5.**
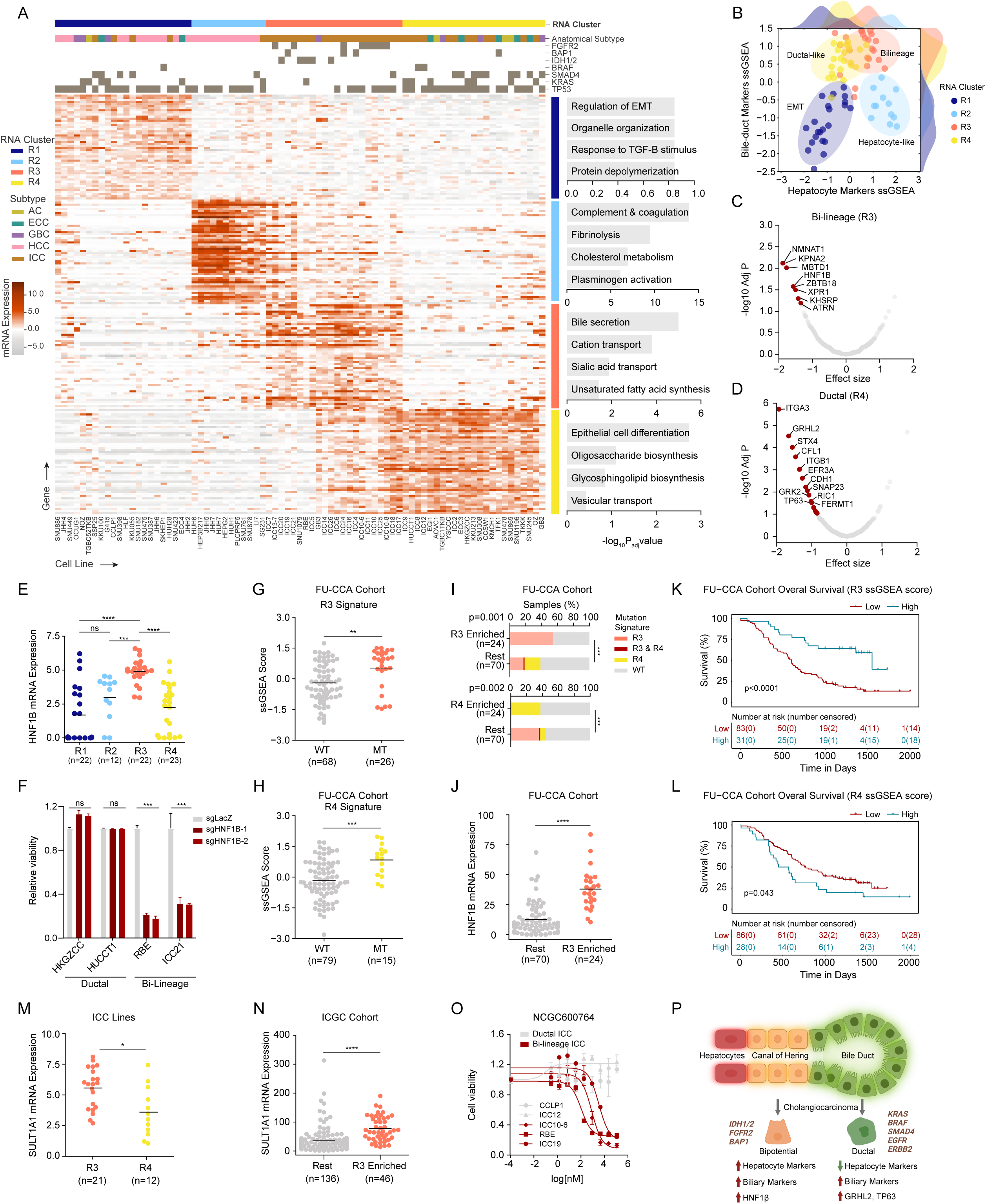
Transcriptomic stratification identifies unique lineage-specific features and vulnerabilities. (A) Heatmap showing normalized mRNA expressions corresponding to the unsupervised clustering of BTC and HCC cell lines based on transcriptomic data. The heatmap displays the top 50 genes enriched in each cluster. Pathway enrichment analysis using the enriched genes for each cluster is indicated on the right. Wilcoxon rank-sum test; adjusted p<0.1. (B) Joint plot showing grouping of cell lines based on hepatocyte and bile-duct markers scoring. Cell lines were given a hepatocyte score (x-axis) and a biliary score (y-axis) based on ssGSEA scoring of hepatobiliary lineage-specific genes. RNA cluster R1 corresponds to EMT subtype, R2-Hepatocyte-like, R3- Bi-lineage and R4-Ductal. (C-D) Volcano plot showing differential dependencies enriched in Bi-lineage (R3) (C) and Ductal (R4) (D) BTC cell lines. Significant genes with at least 2 dependent cell lines in the in-group are highlighted in Red. Unpaired t-test; adjusted p < 0.1. (E) Swarm plot showing HNF1B expression across RNA clusters. Horizontal line segments indicate the mean. Wilcoxon rank-sum; ns, not significant; ***p<0.001, ****p<0.0001 (F) Bar plots showing relative viability after HNF1B knockout in ductal ICC cell lines HUCCT1 and HKGZCC and Bi-lineage ICC cell lines RBE and ICC21. Student’s t-test. ***p<0.001, ****p<0.001 (G-H) Swarm plot showing enrichment of R3 transcriptional signature in WT or MT (IDH1/2, FGFR2, BAP1 alterations) patients (G) or R4 transcriptional signature in WT or MT (KRAS, BRAF and SMAD4 alterations) patient in the FU-CCA cohort (H). Horizontal line segments indicate the mean. Wilcoxon rank-sum; ns, not significant; **p<0.01, ***p<0.001 (I) Stacked bar plot showing percentage of R3, R4 or overlapping genotypes in groupings based on R3 signature enrichment (upper quartile) (top) or in groupings based on R4 signature enrichment (upper quartile) in the FU-CCA cohort (bottom). Fisher’s exact test; **p<0.01, ***p<0.001. (J) Swarm plot showing HNF1B expression levels in R3 enriched patients versus Rest in FU- CCA cohort. Horizontal line segments indicate the mean. Wilcoxon rank-sum test. ****p<0.0001. (K-L) Kaplan-Meier curves showing overall survival in FU-CCA patients (n=114) grouped based on enrichment of R3 transcriptional signature (K) or R4 transcriptional signature (L). (M) Swarm plot showing SULT1A1 mRNA expression in Bi-lineage (R3) (n=21) versus ductal ICC (R4) (n=12) BTC cell lines. Horizontal line segments indicate the mean. Wilcoxon rank-sum test; *p<0.05. (N) Swarm plots showing SULT1A1 mRNA expression in R3 enriched (n=46) patients versus Rest (n=136) in the ICGC cohort. Horizontal line segments indicate the mean. Wilcoxon rank-sum test. ****p<0.0001 (O) Dose-response curves of Bi-lineage (red) (n=3) versus ductal (grey) (n=2) BTC cell lines after 72 hours of treatment with NCGC600764 (P) Schematic describing characteristics of bi-lineage (R3) and ductal (R4) cholangiocarcinoma.

To understand the master regulators of these RNA subtypes, we predicted the activity of 275 transcription factors (TFs) across the cell lines (**Supplementary Figure 5d**, based on DoRothEA TF-target interactions^36^). R1 showed activation of the EMT-inducing TFs (Smad2/3 and TWIST1). R2 was defined by TFs involved in hepatocyte differentiation (HNF4A, HNF1A/B), liver metabolism (PPARA, LXR), and WNT signaling (LEF, TCF4). Endodermal (SNAI, FOXA1, FOXA2, GATA6, GATA4), bile duct/squamous (SOX9, GRHL2, ELF3, TP63), and nuclear hormone receptor family TFs (RARG, NR1H4, NR2F1) were activated in R4. As in the RNA analysis, R3 was a hybrid, sharing features with both R2 and R4. Direct comparison of the main BTC subgroups, showed that R3 was distinguished by the specific activation of TFs governing hepatocyte metabolism (most notably HNF1B), hypoxia response and NF-κB signaling and R4 by specific activation of endodermal and steroid hormone receptor TFs.

The identification of distinct transcriptional subgroups of BTC, characterized by the presence or absence of HCC-like features, is noteworthy in light of the emerging concept that hepatobiliary cancers encompass a spectrum between hepatocyte-like and bile duct-like phenotypes^26,37^. To examine the relationship between cell lineage identity and our RNA clusters, we ranked the cell lines based on the expression of bile duct and hepatocyte markers (Methods). R1 had low scores for both lineage programs, consistent with its EMT signature, R2 had a high hepatocyte score, and R4 had a high bile duct score (**Figure 5b**). R3 stood out in its dual expression of both bile duct and hepatocyte markers. These data indicated that BTCs harbor different cell lineage programs, prompting the designations “EMT” (R1), “Hepatocyte-like” (R2), “Bi-lineage” (R3), and “Ductal” (R4).

Integration of the CRISPR screening data with the RNA clusters revealed selective dependencies in each cluster. These included the EMT-inducing integrin, ITGAV, in R1 and the adhesion-related factors, ITGA3 and CDH1 in R4. Significantly, there were prominent dependencies on lineage-associated TFs that were activated in the respective groups: HNF1A in R2, HNF1B in R3/Bi-lineage, and GRHL2 and TP63 in R4/Ductal (**Figure 5c, d and Supplementary Figure 5e**). Dependency on the liver cell fate transcription factor, HNF1B in R3 was associated with both elevated HNF1B mRNA expression and transcriptional activity (**Figure 5e** and **Supplementary Figure 5f, g**). Importantly, validation studies confirmed that HNF1B in knockout strongly impaired the proliferation of Bi-lineage ICC cell lines while not affecting Ductal ICC cell lines (**Figure 5f** and **Supplementary Figure 5h**). Collectively, these data establish that BTC cell lines comprise distinct lineage states associated with dependency on the regulators of this state (‘lineage addiction’)^38^.

### Hepatobiliary lineage signatures correlate with mutational profiles and outcomes in BTC patient tumors

The lineage subtypes we observed in cell lines exhibited interesting parallels with the recently introduced histological classification of cholangiocarcinoma (CCA), which includes both ICC and ECC^39–42^. This classification proposes stratifying CCA into small duct and large duct subtypes.

Although histologic criteria and molecular profiles are not standardized, small duct CCA shares features with HCC and generally correlates with the enrichment of mutations we observed in R3/Bi-lineage cell lines (BAP1, FGFR2, IDH1/2). Conversely, large duct CCA is enriched for KRAS and SMAD4 mutations, as we find in R4/Ductal cell lines (**Supplementary Figure 5b**). Thus, lineage markers coupled with genomic features could provide an objective measure for defining BTC subtypes.

We tested our lineage-based classification in BTC patient samples (ICGC, FU-CCA and CPTAC-iCCA cohorts) by examining the relationship between mutational profiles and transcriptional signatures. Comparison of tumors with and without alterations in these genes (mutant [MT]) versus WT) by ssGSEA demonstrated that the Bi-lineage genotypes (*IDH1, FGFR2, and BAP1*) had elevated R3 transcriptional signatures, and the ductal genotypes (*KRAS, SMAD4, BRAF)* had elevated R4 signatures (**Figure 5g, h and Supplementary Figure 5i, j**). Similar correlations were observed when stratifying patients based on strong enrichment of the R3 and R4 signatures (upper quartile, ‘enriched’ versus ‘rest’; **Figure 5i and Supplementary Figure 5k, l)**. In addition, given its essential functional role as a lineage survival factor in R3/Bi-lineage cell lines, we examined correlations with HNF1B, specifically. Of note, HNF1B mRNA expression was enriched in patient samples with a high R3 signature (**Figure 5h)** and in those classified as small duct CCA (**Supplementary Figure 5m**). Finally, patients with higher R3 signatures exhibited improved overall survival whereas those with higher R4 signatures showed the opposite (**Figure 5k, l** and **Supplementary Figure 5n, o**).

Collectively, these data indicate that distinct hepatobiliary lineage programs and genetic alterations define clinically significant CCA subtypes, suggesting approaches for molecular stratification similar to those used for other GI cancer types^43^.

### Selective Targeting of Bi-lineage CCA

The above data highlight the possibility of developing therapeutic approaches based on lineage subtype. In this regard, we recently identified a class of compounds that are selectively converted to cytotoxic alkylators by the hepatocyte resident phenol sulfotransferase, SULT1A1^44^, which is strongly expressed in Bi-lineage ICC cells (**Figure 5m**) and in R3 subtype patient samples (**Figure 5n** and **Supplementary Figure 5p**). Prompted by the specificity of the cytotoxic mechanism, we conducted a computational analysis of pharmacogenomic databases to identify additional compounds that could potentially be activated by SULT1A1. This analysis was based on correlations between SULT1A1 levels and drug response profiles (see **Methods**). Testing of known (RITA, N-BIC) and predicted (NCGC600764) SULT1A1-activated alkylators revealed strong selective sensitivity of Bi-lineage ICC as compared to Ductal ICC for each compound (**Figure 5o and Supplementary Figure 5q**). These data reinforce the potential of harnessing SULT1A1 expression as a strategy for the selective targeting of Bi-lineage BTC.

### Quantitative proteomics identifies dysregulated protein networks and associated vulnerabilities in BTC cell lines

Since mRNA and protein expression levels show incomplete correlation^45^, protein quantification can reveal additional distinctions in cell states that are not evident based on RNA analysis alone. We conducted quantitative proteomics analysis of our library of BTC (N=58) and HCC (N=17) cell lines. Median absolute deviation normalization followed by unsupervised clustering identified five distinct proteomic subgroups, which showed broad, but incomplete, concordance with the RNA subgroupings **(Figure 6a, Supplementary Figure 6a** and **Table S6).** Protein cluster P1, enriched for expression of cytoskeletal organization and cell motility proteins, contained most of the R1/EMT cell lines, representing HCC and different BTC subtypes. P2 had elevated expression of RNA processing and splicing factors and was enriched for the R2/HCC group. P3 showed signatures for membrane/vesicle organization and was a blend of each RNA group. P4 was enriched for expression of glycosylation pathways and contained mainly Bi-lineage/R3 lines. P5, enriched for mitochondrial and RNA metabolism signatures, contained exclusively cell lines in the ductal/R4 RNA group. Genomic alterations were also spread across clusters, although trends included absence of KRAS mutations in P4 and enrichment of SMAD4 mutations in P5 (**Supplementary Figure 6b**).

**Figure 6.**
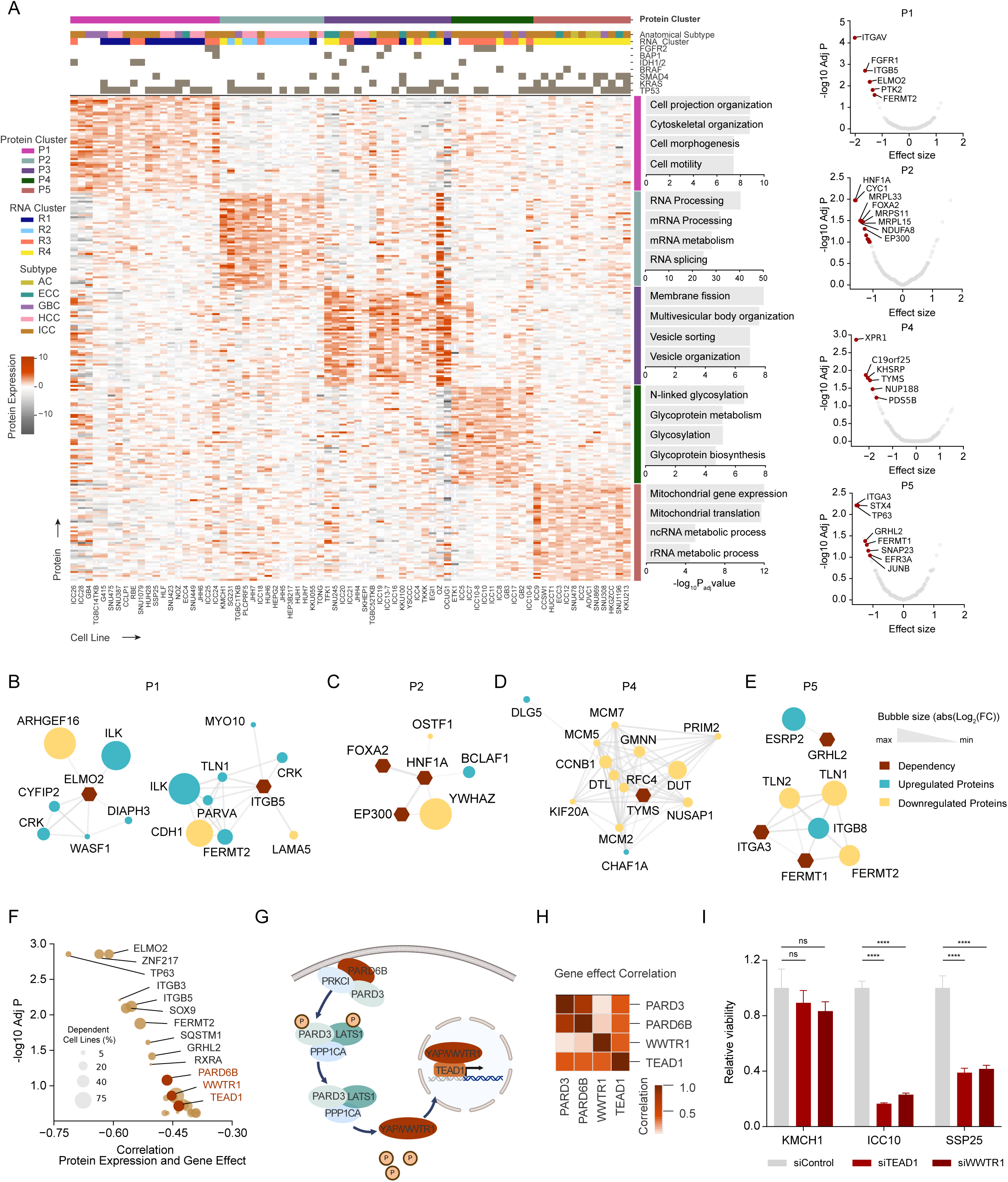
Proteomic characterization of BTC cell lines identifies dysregulated protein networks and associated vulnerabilities. (A) Heatmap showing median-MAD normalized protein expressions corresponding to the unsupervised clustering of cell lines based on protein expression profiles. The heatmap displays the top 50 proteins enriched (Wilcoxon rank-sum, adjusted p<0.1) in each cluster. Pathway enrichment analysis of differentially expressed protein (Methods, adjusted p<0.1) for each cluster is indicated on the right. Each cluster is accompanied by volcano plots indicating differential dependencies corresponding to each cluster (unpaired t-test). significant genes with at least 2 dependent cell lines in the in-group are highlighted in red. Cluster P3 does not exhibit any significant dependencies. (B-E) Protein-protein interaction (PPI) networks generated using STRING database by combining differential dependencies and significantly up and downregulated proteins per cluster. Each network includes first neighbors corresponding to the dependencies. Size of the bubble corresponds to abs (Log_2_ Fold Change) value. Red=Dependency, Blue= upregulated proteins and Yellow= downregulated proteins. (F) Pearson correlation of protein expression and corresponding gene effect scores. Bubble size indicates the percentage of cell lines (n=52) dependent on the specified protein (gene effect<- 0.5). Proteins highlighted in red regulate HIPPO signaling. (G) Schematic representation of PARD3, PARD6B and WWTR1 regulating TEAD1 transcriptional activity. (H) Heatmap showing correlation between PARD3, PARD6B, WWTR1 and TEAD1 gene effect of BTC cell lines (n=39). (I) Bar plots showing relative viability after siRNA mediated knockdown of TEAD1 or WWTR1. Bars represent mean ± SEM. Student’s t-test. ****p<0.0001

We identified top dependencies in these groups and integrated the data with pathway analysis of the differentially expressed proteins. These features were aggregated into interaction networks using the STRING database to define protein markers correlating with dependencies. P1 showed high dependencies on cell motility genes, ITGB5, ELMO2, and FGFR1, consistent with the protein signatures (**Figure 6a (right)** and **Figure 6b**). Cluster P2 was enriched for dependencies on the endodermal TFs, HNF1A and FOXA2, and their binding partner, EP300. (**Figure 6a (right)** and **Figure 6c**). P4 exhibited dependency on the phosphate exporter, XPR1, and on thymidylate synthase (TYMS) (**Figure 6a (right)** and **Figure 6d**). P5 showed dependency on the epithelial differentiation factors, TP63, ITGA3, FERMT1, and GRHL2 (**Figure 6a (right)** and **Figure 6e**). Importantly, in most cases, these dependencies correlated with expression of an interacting protein, suggesting potential biomarkers (e.g. ELMO2/ARHGEF16 and GRHL2/ESRP2; **Supplementary Figure 6c, d**). Thus, proteomics-based clustering identified functional groupings that were not resolved by transcriptomics analysis. Of note, these analyses suggested that BTCs with a ductal RNA signature can be resolved further, with one subgrouping exhibiting increased dependency on an epithelial differentiation program (Cluster P5).

### Upregulation of Hippo pathway proteins predicts dependency

The proteomics data also highlighted specific genetic dependencies that correlated strongly with the expression of their corresponding proteins, with TP63, ZNF217, and ELMO2 showing the most significant associations (**Figure 6f**). Interestingly, protein expression-dependency correlations appeared to be particularly evident for Hippo pathway components. Negative regulation of Hippo pathway kinase, LATS1, by upstream regulators such as the PARD6B/PARD3 polarity complex, leads to induction of the oncogenic WWTR1 and YAP transcriptional coactivators, which promote proliferation in association with TEAD family transcription factors^46–48^ (**Figure 6g**). We found that protein levels of PARD6B, WWTR1, TEAD1, and PARD3 correlated with dependency, whereas only weak correlations were observed for the corresponding mRNA levels (**Supplementary Figure 6e**-g) with the exception of PARD6B).

Dependencies on PARD6B/PARD3 and WWTR1 correlated with TEAD1 dependency (**Figure 6h** and **Supplementary Figure 6h**), highlighting the importance of this protein network. We confirmed that WWTR1 and TEAD1 knockdown suppressed growth of BTC cell lines with high WWTR1/TEAD1 expression (ICC10, SSP25) but not in the low-expressing KMCH1 model (**Figure 6i** and **Supplementary Figure 6i**). TEAD1 is an emerging drug target, and the data suggest the use of this protein network as a biomarker for patient stratification.

### Global analysis of dependencies data identifies functionally related BTC subtypes

Finally, we investigated functional relationships among BTCs, focusing specifically on dependencies, in order to gain new insights into the circuitry of these cancers. We first identified ‘outliers’—*i.e.* dependencies that were most selective and variable within the BTC dataset. This was achieved by performing a normal likelihood-ratio test (LRT, y-axis) and ranking the genes based on median gene effect score (x-axis) (**Figure 7a** and **Table S7**). The top outlier dependencies included receptor tyrosine kinase/MAPK signaling components, p53/p63 pathway genes, endocytosis and intracellular trafficking factors, and folate metabolism enzymes.

**Figure 7.**
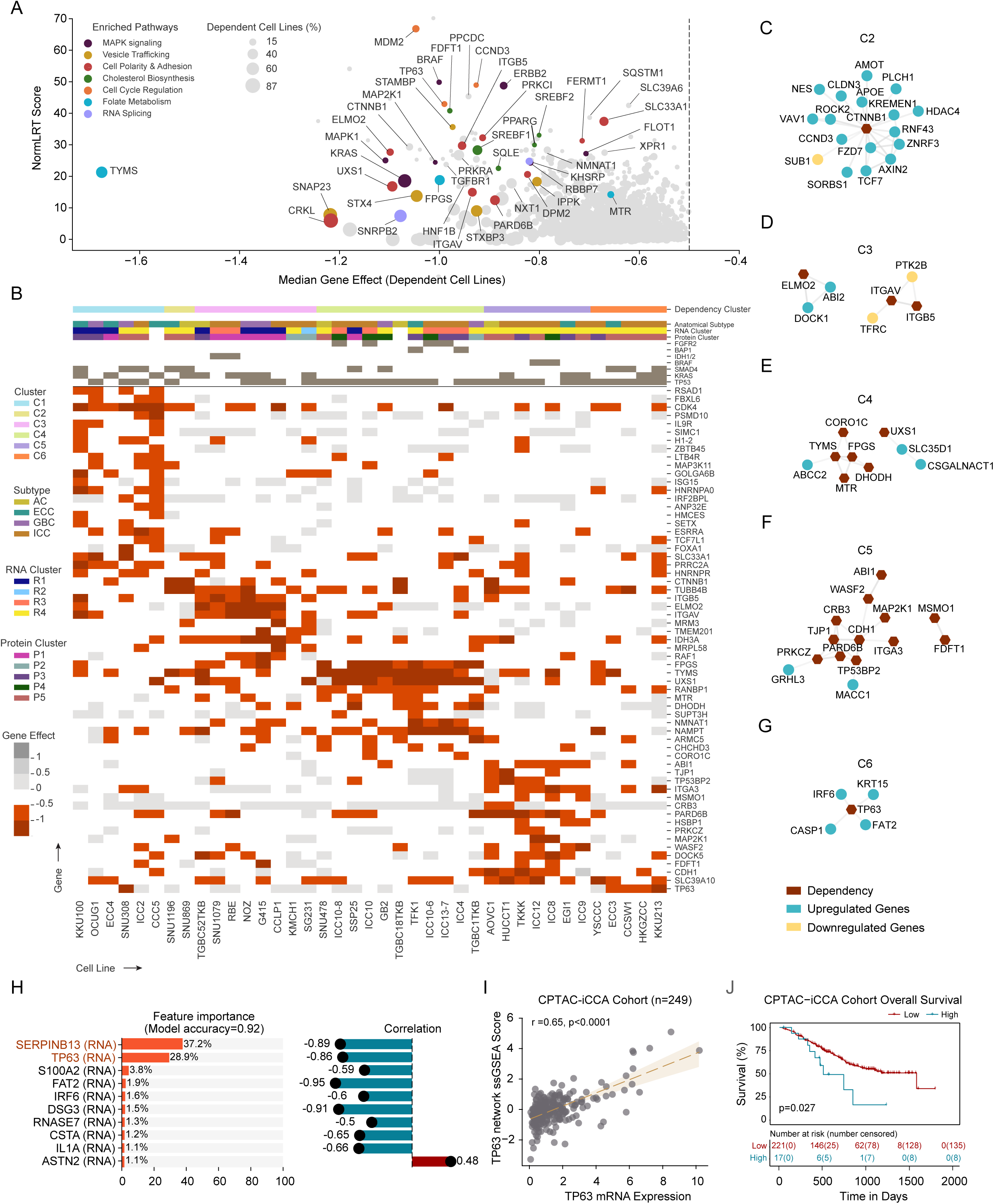
Landscape of genetic dependencies in BTCs. (A) Scatter plot showing outlier dependencies identified by calculating median gene effect scores (x-axis) and normal LRT scores (y-axis) of BTC cell lines (n=39). Colors indicate pathway membership. Bubble size indicates the percentage of BTC cell lines with gene effect score <-0.5. Dotted line indicates the -0.5 dependency threshold. (B) Heatmap showing gene effect scores corresponding to the unsupervised clustering of BTC cell lines based on outlier dependencies. The heatmap displays significant genes with at least 2 dependent cell lines in the in-group. Unpaired t-test; *adjusted p <0.1. (C-G) Protein-protein interaction (PPI) networks generated using STRING database by combining differential dependencies and significantly up and downregulated mRNA features per cluster. Each network includes first neighbors corresponding to the dependencies. Red=Dependency, Blue= upregulated genes and Yellow= downregulated genes. (H) Random Forest analysis showing the top 10 molecular features predictive of TP63 dependency, along with their Pearson correlation with the TP63 gene effect scores among BTC cell lines. (I) Scatter plot showing correlation between TP63 mRNA expression and ssGSEA score using TP63 dependency associated mRNA signature (TP63 network) in CPTAC-iCCA cohort (n=249). Pearson correlation r; Wald test p with t-distribution. (J) Kaplan-Meier curves showing overall survival in CPTAC-iCCA patients (n=238) grouped based on TP63 mRNA expression.

Unsupervised clustering identified six different groups of cell lines (**Figure 7b**). C1, C3, C4 and C5 displayed multiple dependencies distinct to each cluster, whereas C2 and C6 had few shared dependencies (**Supplementary Figure 7a**). To gain a better understanding of the clusters, we examined the functional relationships between the essential genes with each other, as well as with differentially expressed genes in each cluster. These features were aggregated into interaction networks using the STRING database. We also examined associations between genomic alterations or anatomic subtypes of BTC with the clusters (**Supplementary Figure 7b**).

Cluster C1 cell lines (mainly non-ICCs) showed prominent CDK4 dependence (**Figure 7b**). C2 was defined by a network centered on CTNNB1 dependency and characterized by overexpression of WNT/β-catenin pathway target genes and cell junction/polarity factors (**Figure 7c** and **Supplementary Figure 7c**). C3 cell lines exhibited dependency on integrins ITGAV and ITGB5 and on the motility factor, ELMO2, correlating with cell adhesion/migration related expression signatures (**Figure 7d** and **Supplementary Figure 7d**).

Cluster C4 demonstrated selective dependence on nucleotide biosynthesis enzymes (DHODH, FPGS, TYMS, MTR) (**Figure 7e**). Additionally, this cluster exhibited dependency on UXS1 (UDP-glucuronate decarboxylase 1). UXS1 converts UDP-glucuronic acid into UDP-xylose used for proteoglycan generation. UXS1 dependency correlated with expression of the upstream enzyme, UGDH^49^, in line with recent findings that UXS1 prevents toxic build-up of UDP-glucuronic acid resulting from UGDH upregulation^49^ (**Supplementary Figure 7e**). Notably, UXS1 dependency was particularly strong in BTC compared to other cancer cell lines **(Supplementary Figure 7f);** moreover, there were BTC-specific correlations between UXS1 dependency and expression of the nucleotide sugar transporter, SLC35D1 and of CSGALNACT1, which transfers sugars to proteoglycans (**Supplementary Figure 7g, h**). C4 contained the BAP1 and FGFR2 altered cell lines. Accordingly, analysis of patient tumor data demonstrated correlations between these genotypes and increased CSGALNACT1 expression, supporting the relevance of this grouping (**Supplementary Figure 7i, j**).

Top outlier dependencies in C5 produced two functional networks: 1) cholesterol synthesis (MSMO1 and FDFT1), and 2) adhesion, polarity, and cytoskeletal organization (including ITGA3 and PARD6B, and SCAR/WAVE actin polymerization complex^50^ members, ABI1, and WASF2) (**Figure 7f**). WAVE complex dependency correlated with low expression of the ABI1 paralog, ABI2 (**Supplementary Figure 7k, l**), and PARD6B dependency correlated with elevated expression of the EMT-inducing transcription regulator MACC1 (**Supplementary Figure 7m**).

The essentiality of the cellular morphogenesis machinery in this cluster is consistent with the enrichment of MAPK pathway mutations (6/7 lines, including both BRAF V600E models), which potently induce cellular remodeling^51^.

Finally, C6 was defined by pronounced dependency on the driver of squamous differentiation, TP63 (**Figure 7g**), which correlated with elevated expression of TP63 itself and that of a network of squamous markers, including FAT2 protocadherin and SERPINB13 serine protease inhibitor (**Figure 7h** and **Supplementary Figure 7n**). Adenosquamous histopathology denotes a rare and poorly characterized BTC variant. Notably, two of five models in this cluster originated from tumors with adenosquamous carcinomas (ECC3, KKU213). Interestingly, analysis of transcriptomic data of patient tumors showed strong correlation between TP63 expression and the TP63 network observed in C6, which was identified in 4-8% of BTCs (ICGC and CPTAC-iCCA data sets) (**Figure 7i** and **Supplementary Figure 7o**). Of note, higher TP63 expression was associated with poorer prognosis in the CPTAC-iCCA cohort (HR: 1.14, p value=0.07) (**Figure 7j** and **Supplementary Table S7**). Thus, whereas documented adenosquamous histopathology is very rare (<2%), at least partial squamous differentiation driven by p63 appears considerably more common. Across many cancer types, the squamous program of lineage plasticity causes important shifts in drug responses and resistance, suggesting the potential use of expression of the TP63 network in patient stratification.

Taken together, these data elucidate multiple discrete, functionally related BTC subtypes. Although the dependency profiles associated with certain genomic alterations cluster together, the groupings overall transcend individual genotype. These findings suggest opportunities to target convergent pathways across genetic subsets of BTC. They also suggest that factors other than the signature genomic lesions contribute to generating biological heterogeneity in this disease. Notable new targets that may be particularly promising for therapeutic development specific BTC subsets including UXS1, TP63, and the ITGAV/ITGB5 heterodimer^52,53^.

## Discussion

In this study, we present an integrative, molecular and functional genomic analysis of a collection of BTC cell lines, serving as a resource to study this heterogenous group of aggressive malignancies. The atlas comprises a diversity of models, including those newly derived from patients, addressing the overall scarcity of BTC cell lines and the deficiency in key genotypes (e.g. FGFR2, IDH1/IDH2, and BRAF alterations). Incorporating the results into DepMap permitted the identification of broadly shared molecular features and vulnerabilities across BTC lines, distinguishing them from other cancers, while also revealing important biological relationships within BTCs (**Figure 8** and **Table S8**). We defined therapeutic targets associated with predictive markers, including single gene mutations and new, functionally related subclasses of BTCs that extend beyond individual mutations. Importantly, the transcriptional profiles of the cell lines in general, as well as the signatures defining these subclasses in particular, were corroborated in patient samples. These findings support the translational relevance of our results from *in vitro* models to clinical contexts. Thus, the BTC atlas provides biological insights and treatment hypotheses, establishing a robust platform for ongoing research into this tumor type.

**Figure 8.**
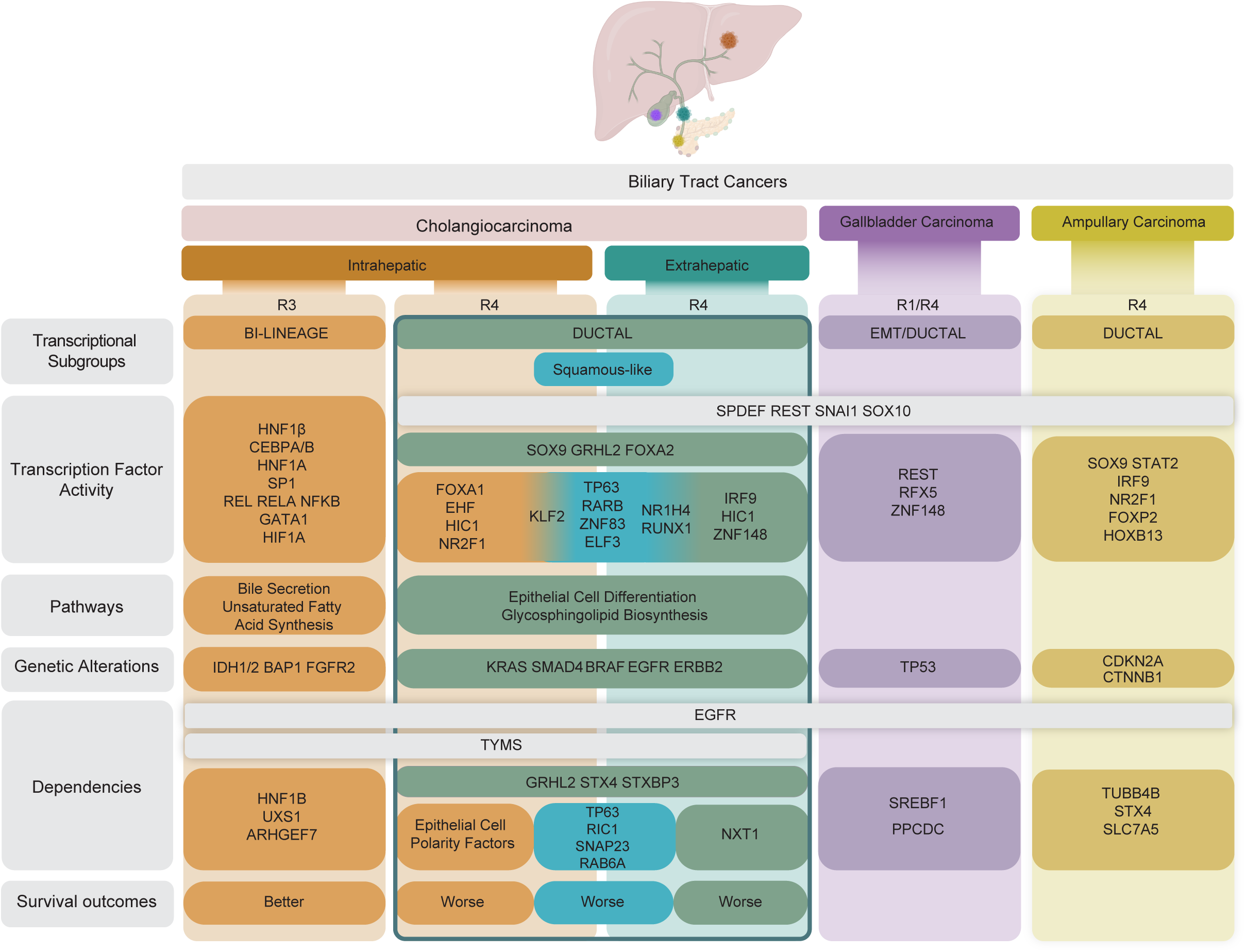
Identification of functional subtypes of BTC. Schematic overview of functional subgrouping of BTC subtypes based on transcriptomic, genomic and dependency profiles.Survival outcomes based on retrospective subgrouping of BTC patient data is indicated where applicable.R1-R4 represents RNA clusters.

Activating genomic alterations in RTK-RAS/PI3K signaling components were common in the cell line collection (EGFR, ERBB2, FGFR2, KRAS, PIK3CA) and typically conferred marked genetic dependencies. The oncogene addiction phenotypes are consistent with the observed clinical efficacy of pharmacological inhibition of these pathways in patients with BTC. We also found that BTCs as a group had a heightened dependency specifically on EGFR compared to cell lines from other tumor types, which, in most cases, was not associated with direct genomic alterations in the pathway. In these cases, expression of LCN2 and MAP3K8, and deletion of chromosome 3p were predictors of EGFR dependency. Moreover, targeting the EGFR/ERBB pathway potentiated the efficacy of mutant KRAS^G12D^ inhibition, overcoming signaling feedback as we previously showed in the context of FGFR inhibition in FGFR2-fusion+ BTC cells. Collectively, these data point to a broad reliance of biliary lineages on the EGFR pathway for cell growth and survival. Whereas EGFR targeting has shown limited promise in unselected patients with BTC, our data support the use of EGFR inhibition in more defined populations (e.g. 3p deletions) and the exploration of EGFR inhibition in combination treatments. Finally, we found that inhibition of PTPN11/SHP2 was highly effective in all FGFR2 fusion-positive models tested—shutting off the MEK/ERK pathway and blocking growth of both cell lines that were sensitive and resistant to FGFR TKI treatment. These observations highlight the potential of SHP2 inhibition to combat polyclonal resistance to FGFR TKI treatment.

For many malignancies, the stratification into tumor subtypes is utilized to guide therapeutic decisions and aid in the interpretation of clinical responses. However, such systems are not currently used in the management of BTC. Whereas WHO has designated histologic subtypes of CCA^54,55^ — small and large duct — the characterization, based largely on duct size, is prone to significant inter-observer variability and is not robust, making it difficult to apply consistently. Thus, there remains the need for a robust classification system linked to disease biology. In this regard, the atlas suggests new opportunities for tumor classification. Through clustering of cell lines based on their transcriptomic profiles, two distinct CCA subtypes were identified, one expressing both hepatocyte and bile duct markers (Bi-lineage) and the other, only bile duct lineage markers (ductal) (**Figure 5** and **8**). These subtypes were characterized by differences in activity of cell lineage transcription factors, genetic dependencies and profiles of gene mutations. Analysis of bulk transcriptomic data from CCA patients revealed that the lineage signatures were enriched for the same gene mutation profiles, and had independent prognostic significance, serving as markers of better (Bi-lineage CCA) and worse outcome (biliary CCA) following surgical resection. It is notable that the dichotomy in lineage features and genomic alterations defining these subtypes is broadly reminiscent of the small ductal/large duct classification, while our studies provide molecular marker profiles and biological insights.

Overall, our cell line data suggest the existence of two major lineage-associated ICC subtypes, driven by distinct hard-wired, cell-autonomous transcriptional programs, and differing in their clinical behavior. Moreover, the dependencies data emphasize the functional relatedness of these subtypes, suggesting opportunities to target vulnerabilities that transcend individual genotypes.

Our data also indicate that a molecular signature for ‘partial squamous’ differentiation designates an additional BTC subtype (**Figure 8**). Clustering of cell lines based on dependencies revealed a group that is highly dependent on p63 for viability, associated with upregulation of select p63 targets. Analysis of patient samples identified this expression signature in 4-8% of patient samples, compared to the exceptional rarity of BTC designated as having squamous histopathology (ref). The partial squamous signature correlated with poor outcomes, consistent with squamous differentiation conferring cellular plasticity and driving treatment resistance in other cancer contexts. The identification of this signature in a significant subset of BTCs suggests an additional approach to patient stratification.

The cell line collection had less representation of BTCs outside of ICC, although we also observed enriched dependencies in the different anatomic subtypes of BTC. Notable findings included the essential roles of PPCDC and SREBF1 in a subset of GBC cell lines. These data are particularly striking considering the overall dispensability of these genes across solid tumor cell lines. PPCDC dependency in GBC is notable in light of the emerging role of increased demand on CoA synthesis in PI3K- and MYC-driven breast cancer^56,57^. The essential role of CoA in fatty acid and cholesterol synthesis suggests a possible link with the parallel dependency on SREBF1 in GBC cells. In ECC, there was a prominent enrichment in dependence on the NXT1 mRNA nuclear export factor. This was associated with low relative expression of its paralog NXT2, indicating that a ’collateral lethality’ relationship contributes to this dependence.

In summary, by integrating molecular profiles and functional annotations, the BTC cell line atlas enhances our understanding of BTC heterogeneity and opens avenues for targeted therapeutic approaches. These studies highlight new BTC subtypes with a shared molecular circuitry while also reinforcing and extending the view that there is marked overall variation in the oncogenic program supporting the growth of different BTC subsets. This biological diversity raises the practical question of how findings from the atlas can be translated to the clinic. Due to their relative rarity compared to other major GI tumor types, there is a challenge in developing tailored clinical trials for the multiple molecular subtypes of BTC. In this regard, innovations such as the development of umbrella trials with a common protocol enable the efficient initiation of investigator-initiated studies across multiple institutions, supported by centralized data monitoring. This approach enhances enrollment and facilitates investigations involving less common disease populations.

## Limitations of the study

It will be important to further expand the atlas to fully capture the heterogeneity of this group of cancers. While we have amassed a substantial collection of BTC cell lines — with the number of models in the DepMap collection comparable to or in some cases exceeding those from more common cancers such as HCC and prostate cancer— additional models are necessary to comprehensively delineate the landscape of vulnerabilities related to genotype, anatomic subtype, and other characteristics. In this regard, the significant variations in BTC prevalence and genomics associated with geography and risk factors underscore the need for ongoing development and analysis of representative sets of models. Examples include liver fluke-associated cholangiocarcinoma^8^ and GBC from regions with notably high incidence rates^58^.

Another limitation of our study is that the CRISPR screen was performed in vitro, which does not fully mimic the growth conditions of tumors in situ. Thus, it will be necessary to validate dependencies in vivo. Moreover, it will also be crucial to continue to correlate features identified in the BTC atlas with primary patient samples. The gene expression-based associations that we present, such as our analysis of the significance of the R3 and R4 lineage groupings for CCA, and partial squamous ICCs, rely on mRNA-seq of bulk tumors. Integrating data from the atlas with spatial transcriptomics and/or single cell analysis of human tumors will be an important next step in elucidating the molecular circuitry of BTC tumor cells themselves and associating this circuitry with functional dependencies and other features.

Overall, further interrogation and expansion of this atlas, in conjunction with analysis of primary human tumors using refined methods, offers promise for advancing understanding and developing therapies for this challenging disease.

## Author contributions

Conceptualization: V.V., L.S., N.B.

Data curation: V.V., N.K., L.S., Y-H.H., P.V., J.M., R.A., H.E., F.V., J.K., M.K., X.I.H.L., R.M.,

W.H. C.R.F., G.G., N.B.

Formal Analysis: V.V., N.K., Y-H.H., R.M., A.P., W.H., G.G.

Funding acquisition: N.B, F.V.

Investigation: V.V., L.S., P.V., J.M., Q.W., Y.Z., P.K., L.K., L.L., I.F., I.G., W.S., H.K., Q.X., E.R.B, M.J.W, S.C.

Methodology: P.V., J.M., R.A., F.V., W.R.S., M.S.L., W.H., G.G.

Project administration: R.K.K., F.V., W.R.S., D.J., W.H., C.R.F, G.G., N.B.

Resources: R.K.K., F.V., W.R.S., D.J., C.R.F., G.G., N.B.

Software: V.V., N.K., Y-H.H., A.P., G.G.

Supervision: V.V., G.G., N.B.

Visualization: V.V., N.K., Y-H.H, L.S., G.G., N.B.

Writing – original draft: V.V., N.B.

Writing – review & editing: N.K., L.S., H.E., R.K.K., J.M.C., V.D., C.R.H., F.V., G.G.

## KEY RESOURCES TABLE

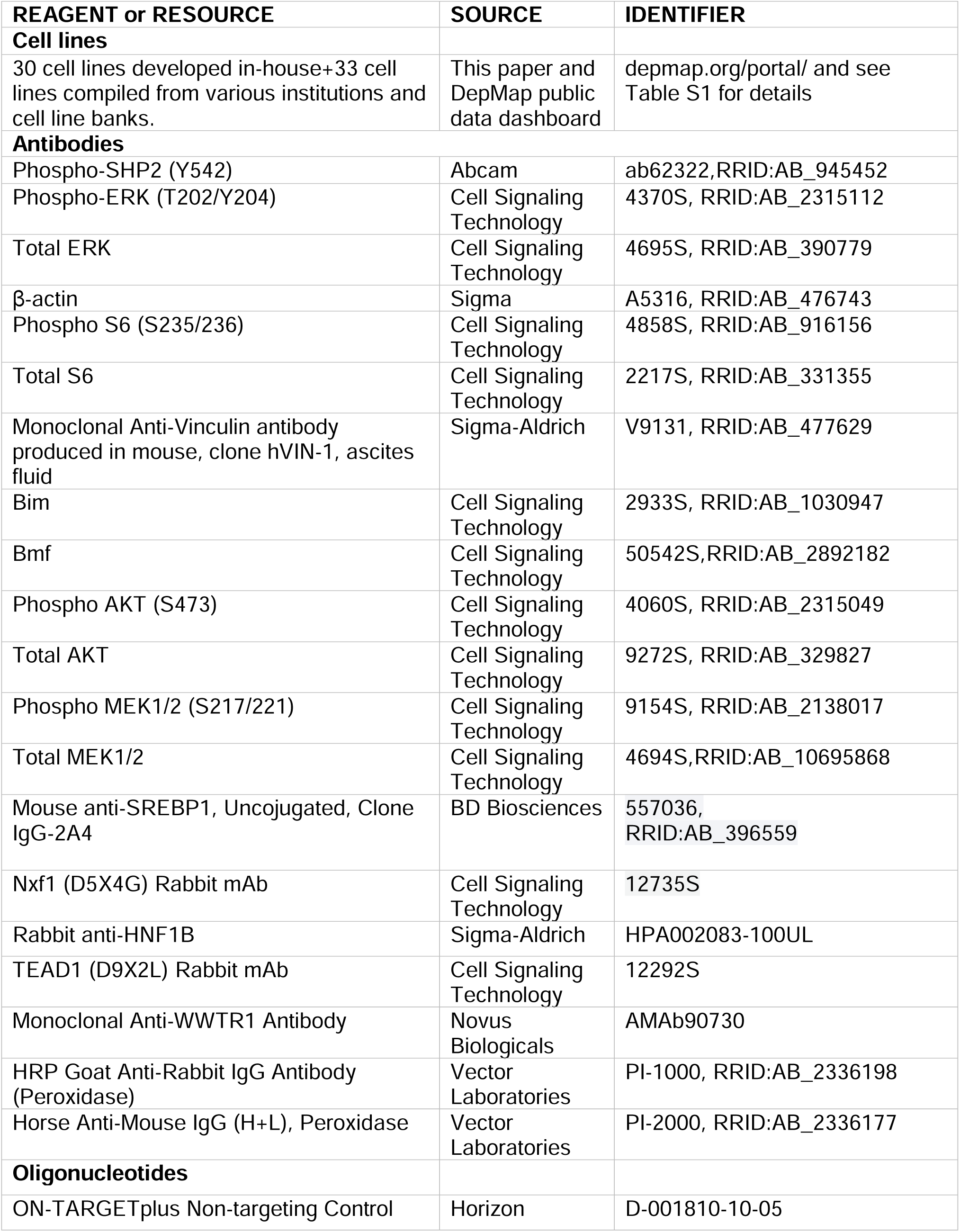
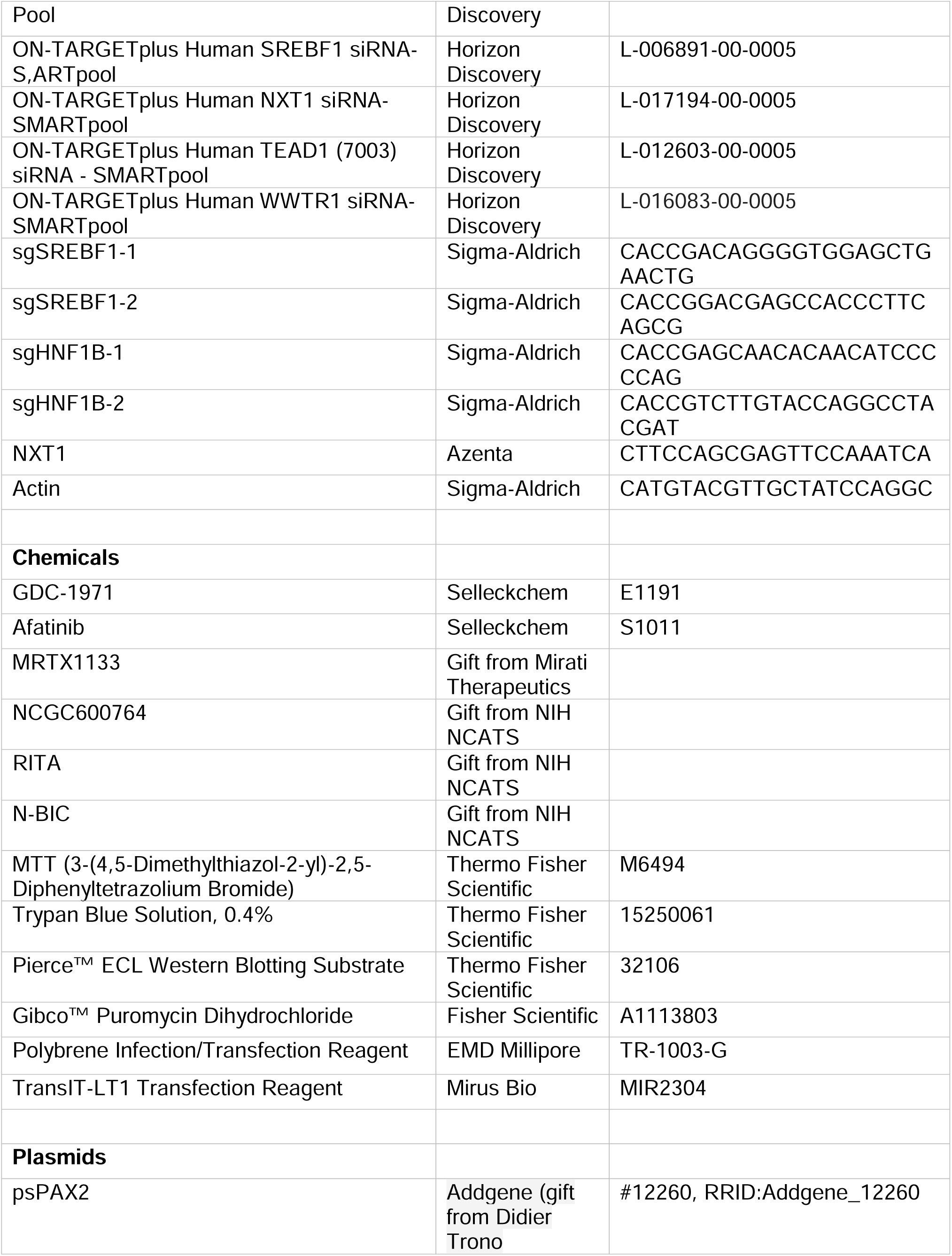
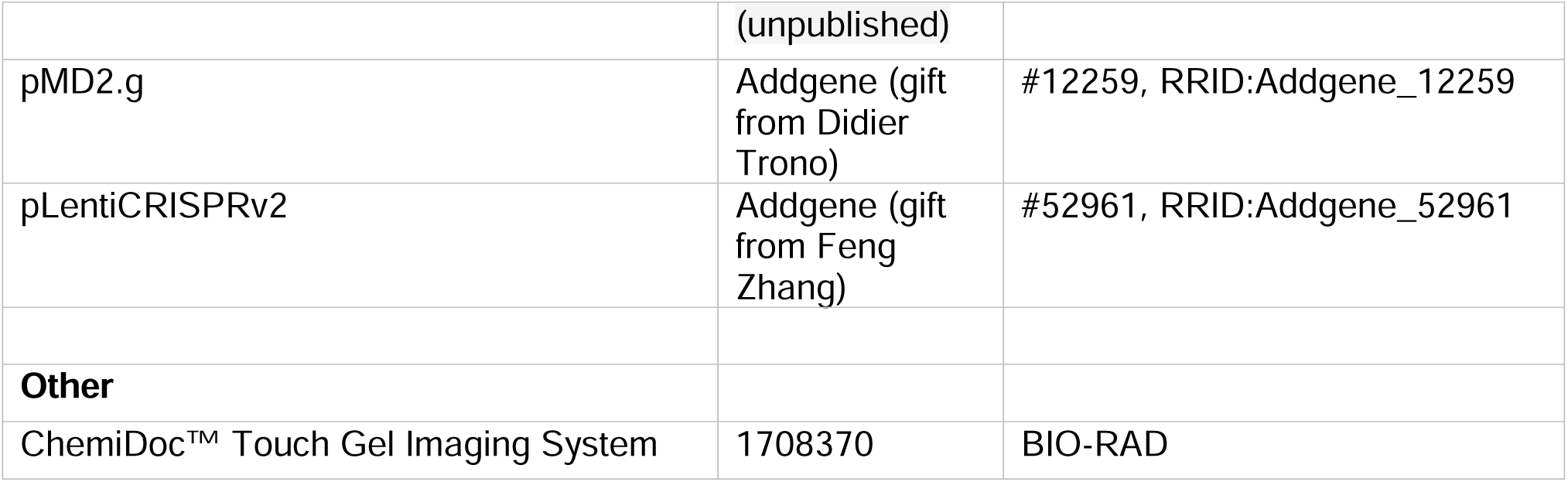

## RESOURCE AVAILABILITY

Requests for reagents and other inquires resources should be directed to the Lead Contact, Dr. Nabeel Bardeesy

### Materials Availability

Cell line, PDX models and reagents used in this study will be made available upon request and with a completed Materials Transfer Agreement.

## EXPERIMENTAL MODELS AND CLINICAL INFORMATION

### Ethics statement

Patient specimens used for cell line development were approved by the Office for Human Research Studies at Dana-Farber/Harvard Cancer Center (protocols 19-699, 14-046 and 02- 240). Diagnostic information from patients are available in Table S1. Animal studies were approved under the protocol 2019N000116 by the Massachusetts General Hospital (MGH) Institutional Animal Care and Use Committee.

## METHOD DETAILS

### Generation of patient-derived BTC cell lines

Cell lines that were developed in-house were established either directly from primary tumor tissues or after at least 3 passages of primary tumors as xenografts in NSG mice. Briefly, resected or autopsy specimens either directly or after growth as a PDX was minced with sterile razor blades, digested with trypsin for 30 minutes at 37°C, and then resuspended in RPMI supplemented with 20% fetal bovine serum (FBS), 1% L-glutamine (Gibco, #25030-081), 1% MEM Non-Essential Amino Acids Solution (Gibco, #11140-050), 1% Sodium Pyruvate (Gibco, #11360-070), 0.5% penicillin/streptomycin (PS), 10 μg/mL gentamicin (Gibco, #15710-064), and 0.2 Units/mL human recombinant insulin (Gibco, #12585-014), seeded on plates coated with rat tail collagen (BD Biosciences), and incubated at 37°C with 5% CO2. All cell lines were adapted to grow in uncoated tissue culture plates in RPMI supplemented with 10% FBS and 1% PS. Cell lines were authenticated by short tandem repeat (STR) profiling at Broad between January and December 2018 and were routinely tested for mycoplasma using LookOut Mycoplasma PCR Kit (Sigma)

### Whole exome sequencing

DNA library preparations, sequencing reactions and bioinformatic analysis were conducted either by DepMap at Broad Institute of MIT and Harvard or Azenta Life Sciences, LLC. (South Plainfield, NJ, USA).

#### Library Preparation and Sequencing

The DNeasy Blood & Tissue Kit (Qiagen) was used for Genomic DNA isolation. After quantification with the Qubit 2.0 Fluorometer (ThermoFisher Scientific, Waltham, MA, USA), libraries were prepared using the Twist Human Core Exome library preparation kit (Twist Biosciences, South San Francisco, CA, USA). In brief, acoustic shearing of DNA was performed using the Covaris S220 instrument. Fragmented DNA was then end-paired and subjected to 3’ end adenylation and adapter ligation. DNA fragments were subsequently enriched by low cycle PCR. Adapter-ligated DNA fragments were validated using Agilent TapeStation (Agilent Technologies, Palo Alto, CA, USA), and quantified using Qubit 2.0 Fluorometer. Adapter-ligated DNA fragments were hybridized with biotinylated baits. The hybrid DNAs were captured by streptavidin-coated binding beads. After extensive washing, the captured DNAs were amplified and indexed with Illumina indexing primers. Post-captured DNA libraries were validated using Agilent TapeStation (Agilent, Santa Clara, CA, USA) and quantified using Qubit 2.0 Fluorometer and Real-Time PCR (KAPA Biosystems, Wilmington, MA, USA. The sequencing libraries were multiplexed and clustered onto multiple lanes of a flowcell. After clustering, the flowcell was loaded onto the Illumina HiSeq instrument according to manufacturer’s instructions. The samples were sequenced using a 2x150bp Paired End (PE) configuration. Image analysis and base calling were conducted by the HiSeq Control Software (HCS). Raw sequence data (.bcl files) generated from Illumina HiSeq was converted into fastq files and de-multiplexed using Illumina bcl2fastq 2.17 software. One mis-match was allowed for index sequence identification.

### Filtering and annotation of Azenta WES data

A subset of the data (13 cholangiocarcinoma cell lines) were characterized by whole-exome sequencing and mutation calling using the GENEWIZ platform (Azenta Life Sciences). To remove common polymorphisms and sequencing artifacts, we removed any variant observed in a Panel of Normals (PoN) based on The Cancer Genome Atlas (TCGA) as previously described^59^. Remaining variants were then annotated for their effects on proteins, according to the GENCODE database. In cases where multiple transcribed isoforms meant a mutation had multiple possible effects, the most deleterious possible effect was used (splice-site > nonsense > missense > synonymous > noncoding). Synonymous mutations and noncoding mutations (those outside protein-coding regions) were excluded from final analyses.

### RNA isolation and Transcriptomic sequencing

Total RNA was isolated using RNeasy plus kits (Qiagen). RNA library preparations and sequencing reactions were conducted by DepMap at Broad Institute of MIT and Harvard or Azenta Life Sciences (South Plainfield, NJ, USA) as indicated as follows: Library Preparation with PolyA selection and Illumina Sequencing RNA samples were quantified using Qubit 2.0 Fluorometer (Life Technologies, Carlsbad, CA, USA) and RNA integrity was checked using Agilent TapeStation 4200 (Agilent Technologies, Palo Alto, CA, USA). The RNA sequencing libraries were prepared using the NEBNext Ultra II RNA Library Prep Kit for Illumina using manufacturer’s instructions (New England Biolabs, Ipswich, MA, USA). Briefly, mRNAs were initially enriched with Oligod(T) beads. Enriched mRNAs were fragmented for 15 minutes at 94°C. First strand and second strand cDNA were subsequently synthesized. cDNA fragments were end repaired and adenylated at 3’ends, and universal adapters were ligated to cDNA fragments, followed by index addition and library enrichment by PCR with limited cycles. The sequencing libraries were validated on the Agilent TapeStation (Agilent Technologies, Palo Alto, CA, USA), and quantified by using Qubit 2.0 Fluorometer (ThermoFisher Scientific, Waltham, MA, USA) as well as by quantitative PCR (KAPA Biosystems, Wilmington, MA, USA). The sequencing libraries were clustered on flowcell lanes. After clustering, the flowcell was loaded on the Illumina instrument according to manufacturer’s instructions. The samples were sequenced using a 2x150bp Paired End (PE) configuration. Image analysis and base calling were conducted by the Control software. Raw sequence data (.bcl files) generated from the sequencer were converted into fastq files and de-multiplexed using Illumina’s bcl2fastq 2.17 software. One mismatch was allowed for index sequence identification. Sequencing data from Azenta was subjected to analysis using DepMap pipelines to harmonize the data.

### Protein isolation and proteomic profiling

Frozen cell pellets were lysed by addition of 500 µL Lysis buffer (75mM NaCl, 50mM HEPES pH 8.5, 10mM Sodium pyrophosphate, 10mM Sodium Fluoride, 10mM B-Glycerophosphate, 10mM Sodium Orthovanadate, roche complete mini EDTA free protease inhibitors, 3% SDS, 10mM PMSF). The disulfide bonds were reduced by adding dithiothreitol (DTT) to a final concentration of 5 mM and incubation at 56 °C for 30 min. Followed by adding iodoacetamide to adjust a final concentration of 15 mM and an incubation in the dark at room temperature for 20 min. The reaction was stopped by adding DTT to a final concentration of 5 mM and incubation in the dark at room temperature for 15 minutes. Protein was isolated by adding one part of Trichloroacetic acid to 4 parts of protein solution and incubation for 10 min on ice. The precipitated protein was pelleted by centrifugation (15,000 g, 10 min, 5 °C) and washed twice with prechilled acetone (- 20 °C, 300 µL, 15,000 g, 10 min, 5 °C). The remaining protein pellets were resuspended in 500µL 1 M urea, 50 mM HEPES or EPPS (pH 8.5) and digested overnight at room temperature with 1 µg/µL endoproteinase Lys-C (Wako) followed by a digestion with sequencing-grade trypsin (Promega) at a final concentration of 1 ng/μL 6 h at 37 °C. The digestion was quenched with 1% trifluoroacetic acid (TFA), and peptides were desalted using Sep-Pak C18 or OASIS HLB solid-phase extraction (SPE) cartridges (Waters). The peptide concentration of each sample was determined using a BCA assay (Thermo Scientific).

For labeling with TMT-10plex or TMTpro reagents (Thermo Scientific), 50 µg for TMT and 25 ug for TMTpro or of peptides were dried and resuspended in 50 µL of 200 mM HEPES or EPPS (pH 8.5), 30% acetonitrile (ACN). Labeling was performed by adding 150 μg TMT or 75 ug TMTpro reagent in anhydrous ACN and incubating at room temperature for 1 h. The reaction was stopped by addition of 5% (w/v) hydroxylamine in 200 mM HEPES (pH 8.5) to a final concentration of 0.5% hydroxylamine and incubation at room temperature for 15 min. Samples were acidified with 1% TFA, and samples were combined. The pooled samples were desalted using Sep- Pak C18 or OASIS HLB SPE cartridges. The combined multiplexed samples underwent a prefractionation on an HPLC under basic pH as previously described and fractions were dried^60^.The dried peptides were resuspended in 5 % ACN/5 % formic acid and analyzed in 3-hour runs via reversed phase LC-M2/MS3 on an Orbitrap Fusion, Orbitrap Fusion Lumos or Orbitrap Eclipse mass spectrometer using the Simultaneous Precursor Selection (SPS) supported MS3 method^61,62^ as described previously^63^. MS2 spectra were assigned using a COMET or SEQUEST-based^64,65^ in-house built proteomics analysis platform^66^ using a target-decoy database-based search strategy to assist filtering for a false-discovery rate (FDR) of protein identifications of less than 1 %^67^. Peptides that matched to more than one protein were assigned to that protein containing the largest number of matched redundant peptide sequences following the law of parsimony. TMT reporter ion intensities were extracted from the MS3 spectra, selecting the most intense ion within a 0.003-m/z window centered at the predicted m/z value for each reporter ion, and spectra were used for quantification if the sum of the S/N values of all reporter ions divided by the number of analyzed channels was ≥20 and the isolation specificity for the precursor ion was ≥0.75. Protein intensities were calculated by summing the TMT reporter ions for all peptides assigned to a protein. Intensities were first normalized using a bridge channel (pooled digest of all analyzed samples in an experiment) relative to the median bridge channel intensity across all proteins. In a second normalization step, protein intensities measured for each sample were normalized by the average of the median protein intensities measured across the samples. The data was adjusted by dividing each protein’s profile across samples by its median followed by log2 transformation. Five different TMT experiments analyzed as mentioned above were combined with 12 selected protein profiles published by the Gygi lab (HEP3B217_LIVER_TenPx02, HEPG2_LIVER_TenPx02, HLF_LIVER_TenPx04, HUH1_LIVER_TenPx03, HUH6_LIVER_TenPx20, HUH7_LIVER_TenPx06, JHH4_LIVER_TenPx40, JHH5_LIVER_TenPx04, JHH6_LIVER_TenPx41, JHH7_LIVER_TenPx05, SNU423_LIVER_TenPx42, SNU449_LIVER_TenPx39)^68^. The datasets were combined by rows (proteins) which were normalized to have the same average across all datasets followed by normalization of the columns (samples) such that their median values are adjusted to the average of the medians across all samples.

### Statistical methods

We conducted statistical analysis using both Python and R software tools; corresponding software is explained in the respective methods sections. When examining significant differences between two groups, we employed the Wilcoxon rank-sum test, accompanied by effect size r, with the exception of gene effect scores and growth/viability studies, where we used the unpaired t-test, as per^69,70^ along with Cohen’s d effect size. To adjust for false discovery rates (adjusted p), we utilized the Benjamini-Hochberg method^71^ for multiple testing. A p-value < 0.05 and an adjusted p-value < 0.1 were considered significant, while an adjusted p- value < 0.25 was considered near-significant. Comparison between categorical variables was done using Fisher’s exact test, and to measure correlations between two continuous variables, we employed Pearson correlation. Additionally, for calculating p-values of linear models, the Wald test with a t-distribution was used, with the null hypothesis that the slope is zero. All tests are two-tailed unless otherwise specified.

### Copy number alteration analysis

#### Cross-comparison with patients data

We utilized GISTIC^72^ available on the GenePattern^73^ portal to identify significant copy number alterations in BTC cell lines and the TCGA^74^ datasets. Since the reported segment means of the cell lines are approximately 2 raised to the power of the mean log2 copy ratio, we log-transformed them before inputting them into GISTIC (e.g., log2(x+1)-1).

#### 3p13 arm analysis

We obtained a list of frequently deleted genes for each arm among BTC patients from the TCGA dataset and employed FGSEA using the fgsea package in R to determine the enrichment score of each arm. This score was based on the ranking of the genes, determined using correlation scores of their mRNA expression values and the EGFR gene effect scores.

### Dependency prediction and feature selection

We used the random forest regression model^75^ to predict gene effect scores and rank the features for the 39 BTC cell lines. In order to train the model, we utilized mRNA expression, copy number, fusion, and mutations, which were divided into three categories: hotspot mutation, damaging mutation, and other mutations. Moreover, we considered each BTC subtype as a feature and employed 10-fold cross-validation to train the model. Due to the limited size of our dataset, we enhanced our model’s performance by augmenting the BTC training set with the remaining CCLE dataset (n=985). However, during each iteration of the training process, we only considered the top 1000 features that showed correlation with the target within the BTC training set. We evaluated the accuracy of the model by measuring the Pearson correlation between the predicted gene effect scores and the actual values across all folds. Additionally, we assessed the importance of each feature for predicting each dependency by calculating the average of the scikit-learn’s Random Forest Regressor’s feature importance over the 10 folds.

### Unsupervised clustering

Our approach to clustering mRNA expression, protein expression, transcription factor activities, and the gene effect score datasets involves combining consensus clustering with Louvain clustering as the base model. We utilize the correlation matrix of positive values to identify highly correlated clusters. The generalized Louvain clustering is governed by a parameter gamma, which controls clusters’ resolution. By setting gamma>1, we can achieve greater granularity and smaller clusters. To establish stable clusters, we chose multiple values of gamma. For all datasets except proteomics, we selected a set of gamma parameters [1, 1.5, 2, 2.5, 3, 3.5], and applied clustering 100 times with different seeds, resulting in 600 solutions for each dataset. For the proteomics data, we observed that smaller values of gamma resulted in better identification of data’s inherent modular structure. Hence, we chose a set of gamma parameters [.8, 1, 1.2, 1.4] and applied clustering 200 times for each value, resulting in 800 solutions.

We followed the consensus clustering algorithm as described in^76^ to obtain the final clusters; the implementation is done in Python by incorporating the netneurotools package (https://netneurotools.readthedocs.io/en/latest/).

In summary, to obtain the final clusters, first a cumulative consensus matrix is created that indicates the number of times two points were in the same cluster. Next, it is normalized to obtain a probability distribution. The algorithm then discards the entries below a threshold τ to account for noisy vertices. We followed^77^ to select the parameter τ, where a null model was created by permuting the labels in each solution. Then, the probability distribution of the consensus matrix for the null model was calculated, and the mean of entries was taken as the parameter τ.

The Louvain algorithm was then applied to the new matrix 10 times to obtain a new set of solutions and a new consensus matrix. This process was repeated until the consensus matrix is turned into a block diagonal matrix.

#### Clustering similarities

We used the adjusted Rand index^78^ to assess the agreement between two partitions of cell lines, each consisting of an in-group versus an out-group based on either a cluster or a genomic feature. For this, we utilized the scikit-learn package in Python. A value of 0 indicates that the similarity between the two partitions is no better than random labeling, a value of 1 indicates strong agreement, and negative values, bounded by -0.5, indicate disagreement.

### CRISPR analysis

We used Chronos gene effect scores^79^ and set a threshold of <-0.5 to identify gene dependencies.

#### Exclusion list

We compiled an exclusion list of genes comprising the nonessentials, common essentials (except for KRAS), and inferred common essential genes from the DepMap portal to exclude from certain analyses.

#### Preferential list

We created a list of preferentially essential genes for use in further analyses. First, for each gene, we normalized the gene effect scores using z-score across all CCLE datasets (1,207 cell lines) and then selected the top 20 dependencies for each cell line; this amounted to 696 genes for the BTC dataset. Similarly, we computed the list of preferential genes for 61 Pancreatic Adenocarcinoma cell lines, resulting in 1,045 genes, and for 21 Hepatocellular Carcinoma cell lines, resulting in 388 genes.

#### Co-dependencies

Co-dependencies among BTC cell lines were identified using the Pearson’s correlation of preferential genes across all other genes.

#### NormLRT scores

We applied normality likelihood ratio test (NormLRT) across BTC cell lines using the R implementation of^80^ (https://github.com/ndharia-broad/peddep). NormLRT^81^ is an estimate of how much a distribution of gene effect scores deviates from the normal distribution, therefore, for the subsequent analyses, we only considered genes that scored high, and their distributions were left skewed, indicating a selective dependency.

#### Clustering

We performed the clustering analysis using the 39 BTC cell lines with the CRISPR data. To compute the correlation matrix of cell lines, we selected 417 genes by filtering out the exclusion list and selecting the genes with variance within the top 15%, and NormLRT score within the top 10%. To identify significant features for each cluster, we used t-test and retained significant features with a gene effect score < -0.5 in at least two cell lines. Additionally, we conducted a differential expression analysis using the limma package (further details provided in the corresponding method section) to identify expressed genes associated with each cluster.

We selected genes for downstream analysis using the ’filterByExpr’ function from the EdgeR package, which resulted in 11,919 genes.

#### Differential dependencies

We identified significant dependencies between in-group and out- group classifications for the mRNA (n=58) clusters, protein (n=52) clusters, and across subtypes including ICC, GBC, ECC, AC, and HCC (n=60). To adjust for the inflation observed in p-value QQ plots, we conducted multiple steps to select genes. First, we excluded genes from the exclusion set. Next, we identified the top 100 most variable genes and added genes with a NormLRT score within the top 2%. We also selected genes exhibiting a variance > 0.1 (representing the upper tail of the distribution) from a list of cancer hallmark genes obtained from COSMIC. Finally, we added the list of preferential genes. The union of all these gene sets resulted in 771 genes for the mRNA dataset, 766 genes for the proteomics dataset, and 772 genes for the subtype dataset. We used t-test followed by Benjamini-Hochberg multiple test correction to determine significant dependencies. We also conducted differential dependency analysis for specific genotypes across our BTC cell lines (n=39). We followed a similar methodology for gene selection; however, we adjusted our criteria to include genes within the top 50% of NormLRT scores while being able to effectively control the inflation of p-values and maintain statistical power. This resulted in 5,018 genes, enabling broader exploration of genotypes.

### mRNA expression analysis

#### Normalization

We normalized the log-transformed TPMs using median centering and utilized them in the subsequent analyses, including ssGSEA using the GSVA library in R.

#### Clustering

We used the normalized expression values to generate the correlation matrix of cell lines as input to the clustering algorithm. To identify the signatures of each mRNA cluster, given the variation in gene expression patterns within cluster R3, we opted to utilize the Wilcoxon rank-sum test and selected the top 50 significant features with the largest effect size.

#### Differential expression analysis

We used gene-level expected counts processed by the RSEM software^82^ and used the Limma-Voom package^83^ in R to identify differentially expressed genes.Samples were normalized using trimmed mean of M values (TMM) normalization via ‘calcNormFactors’ function and the Limma package was applied in accordance with standard procedures utilizing ‘lmFit’ and ‘eBayes’ functions to identify differentially expressed genes between in-group and out-group classifications.

#### Pathway enrichment analysis

Pathway enrichment analysis was performed using significant differentially expressed genes (adjusted p<0.1 and log_2_FC<-1.5 or >1.5 fold) identified for each cluster on EnrichR (PMID: 23586463, PMID: 27141961, PMID: 33780170). Gene Ontology terms were used to identify enriched pathways.

### PDX-Cell line comparison

We used normalized mRNA expression values and applied the K-nearest neighbor algorithm with Euclidean distance to find the closest cell line to each PDX. The implementation was done with the ‘NearestNeighbors’ function from the scikit-learn package in Python.

### UMAP visualization

We conducted all UMAP visualizations and preprocessing steps in R. We utilized TPMs and removed genes with combined TPMs less than 10 across both datasets. To compare with TCGA datasets, we utilized RSEM expected counts. For each analysis, we first combined the datasets. We then applied variance stabilizing transformation using the ’vst’ function from the DESeq2 package. Next, we used the Limma package to address the batch effect between the datasets. Finally, we employed the umap package to visualize the two-dimensional projection of the datasets. We conducted this analysis in multiple runs with varying seeds to ensure that the proximity of embeddings is not influenced by randomness.

### Transcription factor activity analysis

We used the DoRothEA R package^36^ (https://saezlab.github.io/dorothea/) to infer transcription factor activities of cell lines. DoRothEA assigns a confidence level (A-E) to each TF-target interaction based on the number of supporting evidence. Hence, we removed the low-confidence (E) TF-target interactions in the subsequent analysis.

We filtered out normalized mRNA expressions for genes with combined expression values less than 45. The remaining genes were used to infer the scored transcription factors using the selected list of TF-target interactions. This resulted in 275 transcription factors and their activity scores for each cell line. Next, we standardized each transcription factor’s activity score across the cell lines using z-score and computed the correlation matrix of transcription factors as input to the clustering procedure.

### Protein expression analysis

We applied the clustering algorithm to 5,299 proteins, which we retained after filtering out the proteins with more than 10% zero or missing values. The selected proteins underwent the median-MAD normalization and were used to calculate the correlation matrix of cell lines. This served as the input for the clustering process. To identify the signatures of each cluster, we performed differential expression analysis (see below).

To perform the remaining analyses, such as correlations, we first filtered out proteins with over 40% zero or missing values, followed by median-MAD normalization.

#### Differential protein analysis

Based on the five defined protein sample clusters we identified differentially expressed proteins between each cluster versus all other clusters. For each comparison a model matrix was generated stratifying samples between the cluster of interest and all other clusters. The linear model fitting on the model matrix followed by empirical Bayes moderated t-test statistic and p-value calculation from the Limma R package (default parameters) was used to determine significant changes in protein levels between the clusters. Protein changes adjusted p-value < 0.1 were considered significant.

#### Functional Enrichment

Functional term enrichment was determined for each set of differentially expressed proteins using the over-representation hypergeometric test “enrich” function from the GSEApy Python module^84^. For each protein cluster the significant up- regulated (adjusted p-value<0.1 & logFC>0) and down-regulated (adjusted p-value<0.1 & logFC<0) proteins were identified and their uniprot protein names were converted to the associated primary gene symbol. The enrichments of the up-regulated and down-regulated genes were calculated separately across the following geneset collections; MSigDB C2 canonical pathway (CP), C5 GO biological process and Hallmark (v2023.1)^85,86^, and EnrichR PPI_Hub_Proteins^87,88^. We used the set of all differential expressions tested protein associated genes as the custom background for this analysis.

#### Protein-protein interaction networks

Protein-protein Interaction (PPI) networks were created using the STRING database and visualized using Cytoscape. Networks were made using dependencies and top features (gene/protein as indicated) identified for each cluster (adjusted p-value<0.1) and filtered for first neighbors, where each bubble denotes a dependency or associated gene/protein, edge denotes confidence of interaction and size of the bubble denotes magnitude of fold change in expression (when applicable).

### Survival analysis

We conducted Kaplan-Meier survival estimates on two groups of patients, stratified using the ssGSEA scores obtained from mRNA clustering signatures. Based on the distribution of scores, we selected the higher quartile as the “High” group enriched in the signatures, and the rest as the “Low” group. We also conducted multivariate cox regression across available clinical data after correcting for multicollinearity, along with the ssGSEA scores. We followed similar steps for the survival analysis based on TP63 mRNA expression values. However, for each dataset, the patients were stratified into low and high groups based on the higher tail of the distribution of TP63 mRNA expression values. All survival analyses were conducted in R using the survival package.

### Dose-response assays

To test drug efficacy, cells were seeded at a density of 2500 cells per well in 96-well plates and incubated overnight to allow attachment. Compounds serially diluted over a 9-point concentration range (as indicated) were added the next day and the cells were incubated for 72- 96 hours depending on the growth rate of individual cell lines. Cell viability was then measured by performing an MTT(3-(4,5-dimethylthiazol-2-yl)-2,5-diphenyltetrazolium bromide) assay. IC_50_ values were calculated using GraphPad Prism 10 by applying a 3-parameter dose-response model.

### Lentivirus production

HEK-293T cells were transfected with a mixture of 6μg of psPAX2 (Addgene), 3µg of pMD2.G (Addgene) and 6µg of vector plasmid in 1.5ml of Opti-MEM (Fisher Scientific) and 45µL of TransIT-LT1 (Mirus Bio) transfection agent in a 10cm tissue culture dish. Virus-containing supernatant was collected at 24 and 48 hours after transfection and either used immediately to infect target cells or aliquoted and stored at -80°C. List of Guide RNA sequences used in this study is provided in the key resources table.

### Immunoblot analysis

Cells were lysed using Pierce^TM^ RIPA buffer (Fisher Scientific) supplemented with 1X Halt Protease and Phosphatase inhibitor cocktail (Thermo Scientific) and the concentration of isolated proteins was quantified using Pierce^TM^ BCA Protein Assay kit (Thermo Scientific). A total of 30µg of protein was used for analysis using SDS-PAGE, electro transfer and blotting with indicated antibodies. Proteins were separated by electrophoresis on 4–20% Mini- PROTEAN® TGX™ Precast Protein Gels (Bio-Rad) and transferred to Immobilon®-P PVDF Membrane (EMD Millipore). PVDF membranes were blocked for 1 hour in 5% Bovine Serum Albumin (BSA) (Fisher Scientific) dissolved in 1X Tris-Buffered Saline with 0.01% Triton-X-100 (TBST) followed by incubation with the primary antibody (1:1000 in blocking buffer) overnight. Following this, membranes were washed in 1X TBST 5 times and incubated with horseradish peroxidase (HRP)-conjugated secondary antibody (1:1000 in 1X TBST) (Vector Laboratories) for 1 hour. The membrane was washed again, and proteins were detected using 1:1 solution of Pierce™ ECL Western Blotting Substrate (Thermo Scientific). A complete list of all primary and secondary antibodies used in the study is provided in the key resources table.

### In vivo tumor growth measurement

All mice were housed in a specific pathogen-free environment at the MGH and treated in strict accordance with protocols 2019N000116 approved by the Subcommittee on Research Animal Care at MGH. For testing in vivo efficacy of SHP2 inhibition, 2 × 10^6^ ICC11 cells were injected subcutaneously into the flank of NSG mice (NOD.Cg-Prkdcscid Il2rgtm1Wjl/SzJ, 00557, The Jackson Laboratory). Tumor-bearing mice (n=6) were randomized into two groups, following which the mice were treated with vehicle (15 w/v% Hydroxypropyl-β-cyclodextrin dissolved in Citrate Buffer, pH 3.0), or GDC-1971 3 mg/kg daily for continuously 28 days by oral gavage. Tumor volumes were monitored by digital calipers. For validating effect of SREBF1 depletion in vivo, 2 × 10^6^ SNU308 cells lentivirally infected with sgLACZ or two distinct guides targeting SREBF1 (sgSREBF1-1 and sgSREBF1-2) were injected subcutaneously into the flank of NSG mice (n=4 mice per cohort). Tumors were measured every 4 days using digital calipers once the tumors in the control group reached a volume of 100mm^3^.

### Quantitative RT-PCR

Total RNA isolated from cells was reverse transcribed using LunaScript® RT SuperMix Kit (New England Biolabs) to generate cDNA as per the manufacturer’s instructions. Quantitative RT-PCR was performed using SYBR® Green Master Mix (Bio-Rad) in a Bio-rad CFX-384 lightcycler (Bio-rad). PCR reactions were run in triplicates and relative expression of target gene was determined using the comparative CT method with actin expression as control. qPCR primer sequences used in this study are listed in the key resources table.

