## Supplementary figures and images for "Generation of a biliary tract cancer cell line atlas reveals molecular subtypes and therapeutic targets"

### Supplementary Figure1.pdf

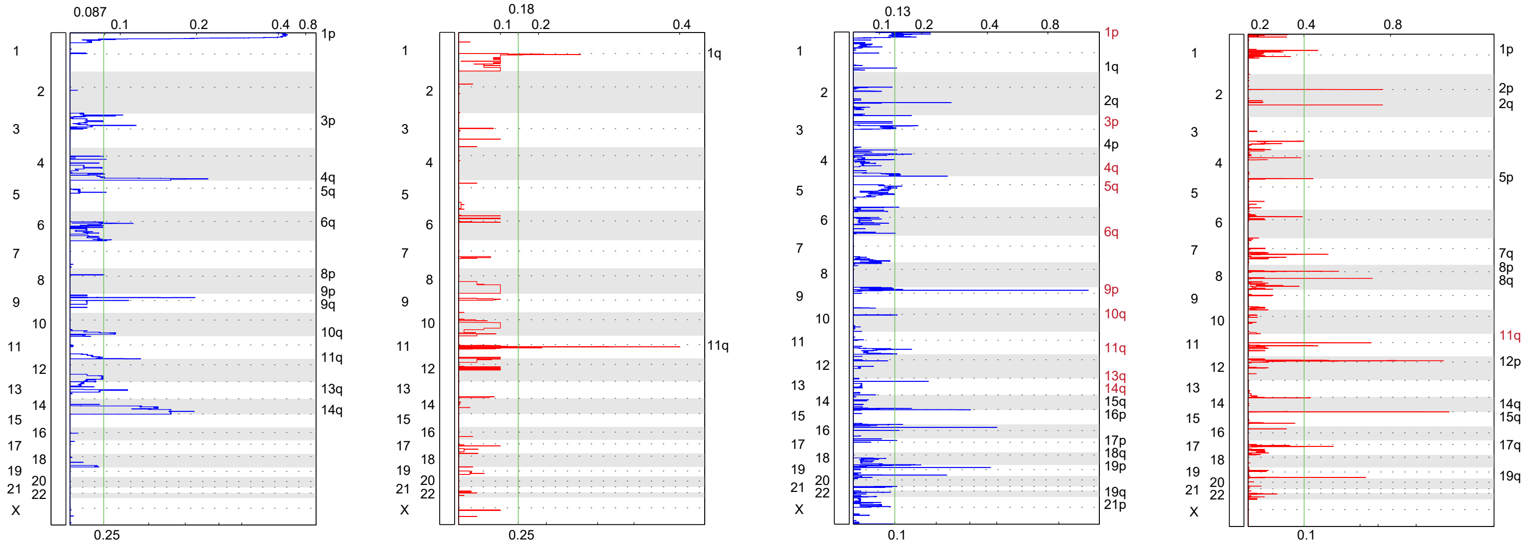

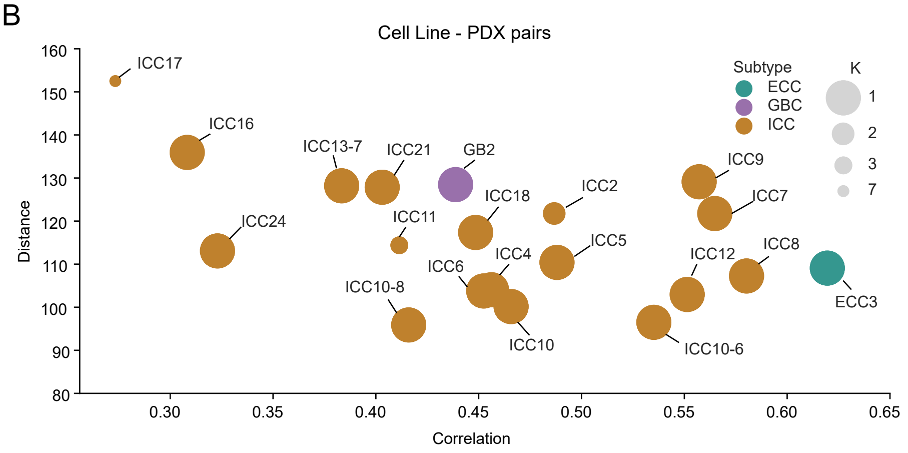

### Supplementary Figure2.pdf

#### Supplementary Figure 2

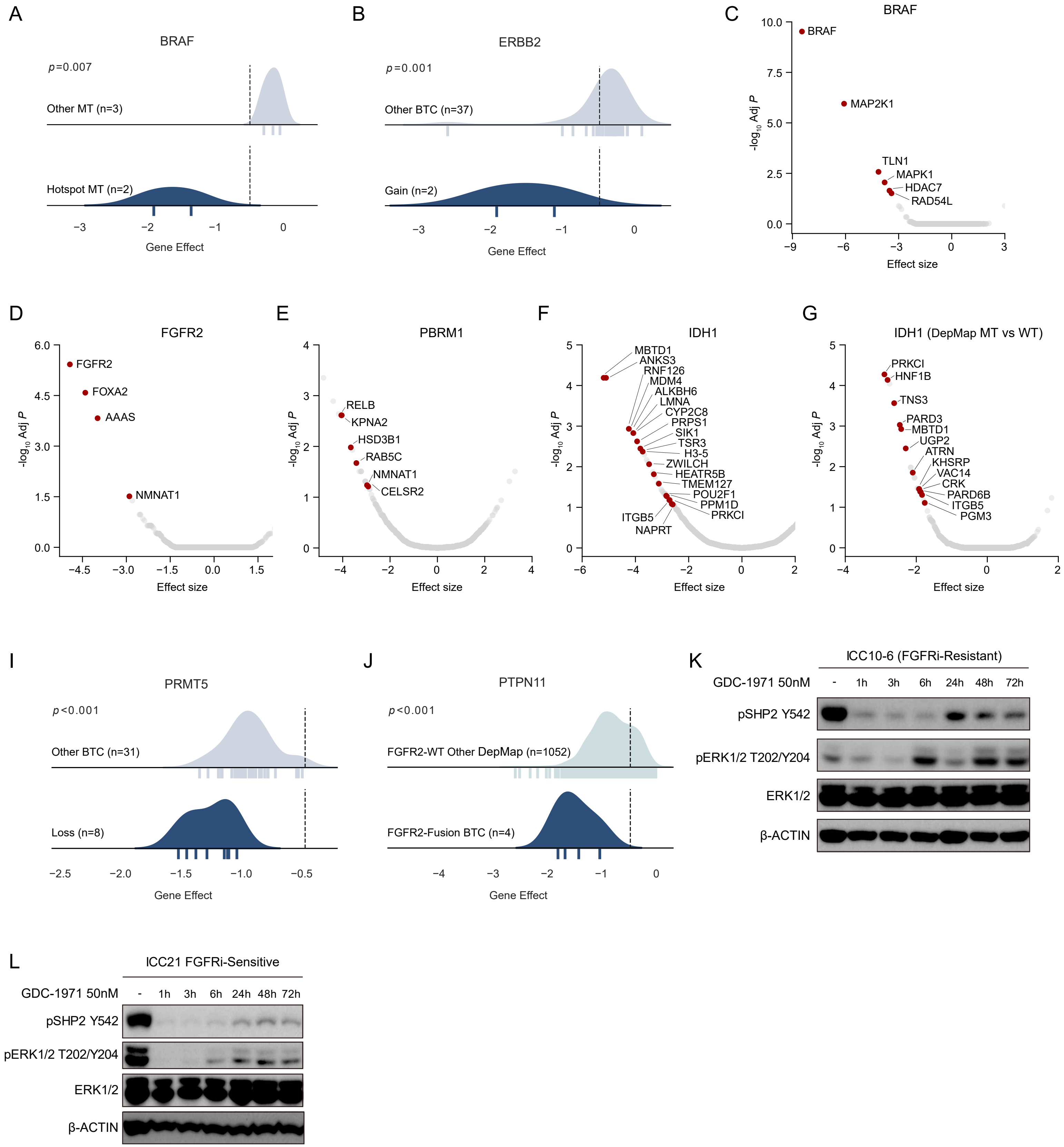

### Supplementary Figure3.pdf

#### Supplementary Figure 3

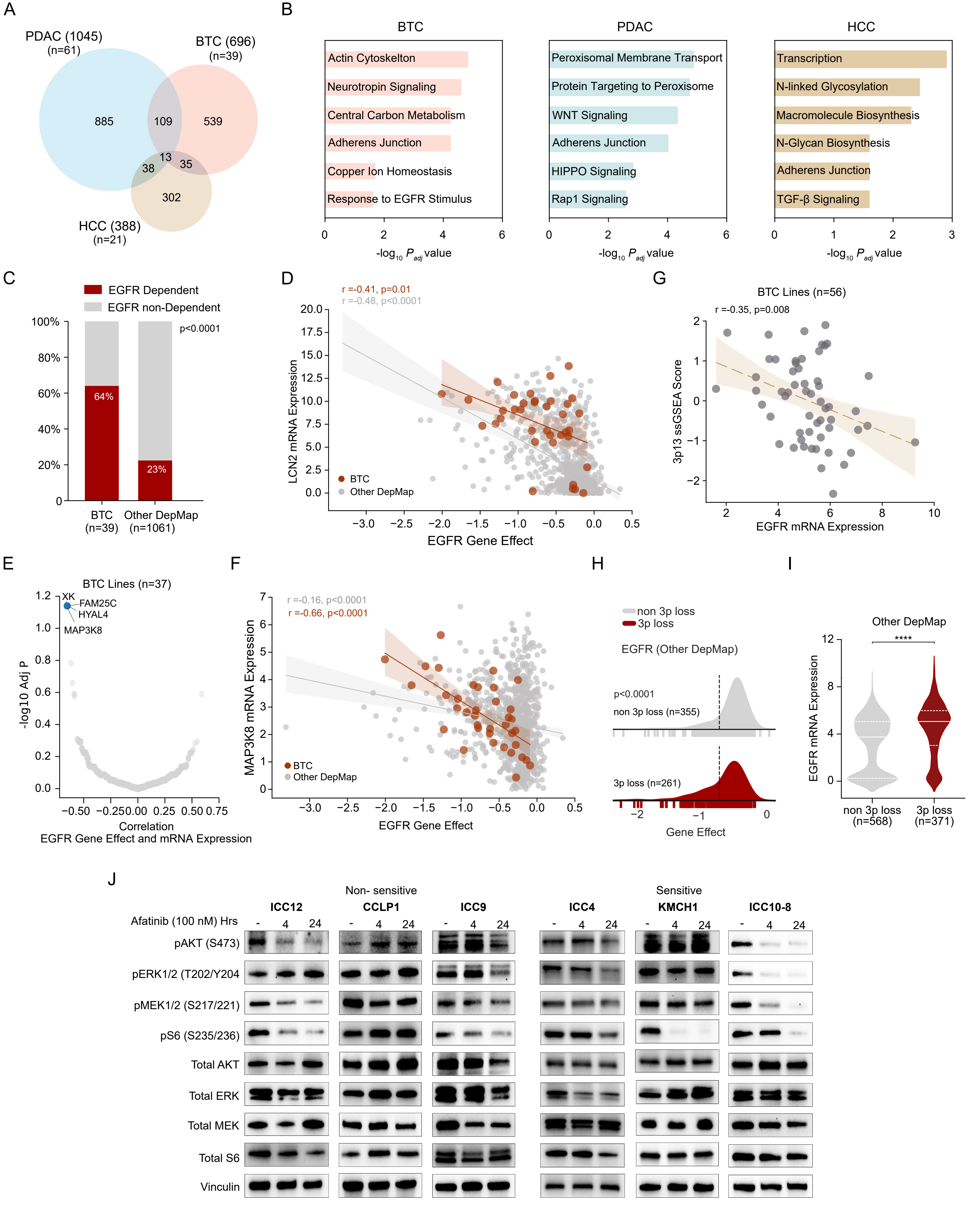

### Supplementary Figure4.pdf

#### Supplementary Figure 4

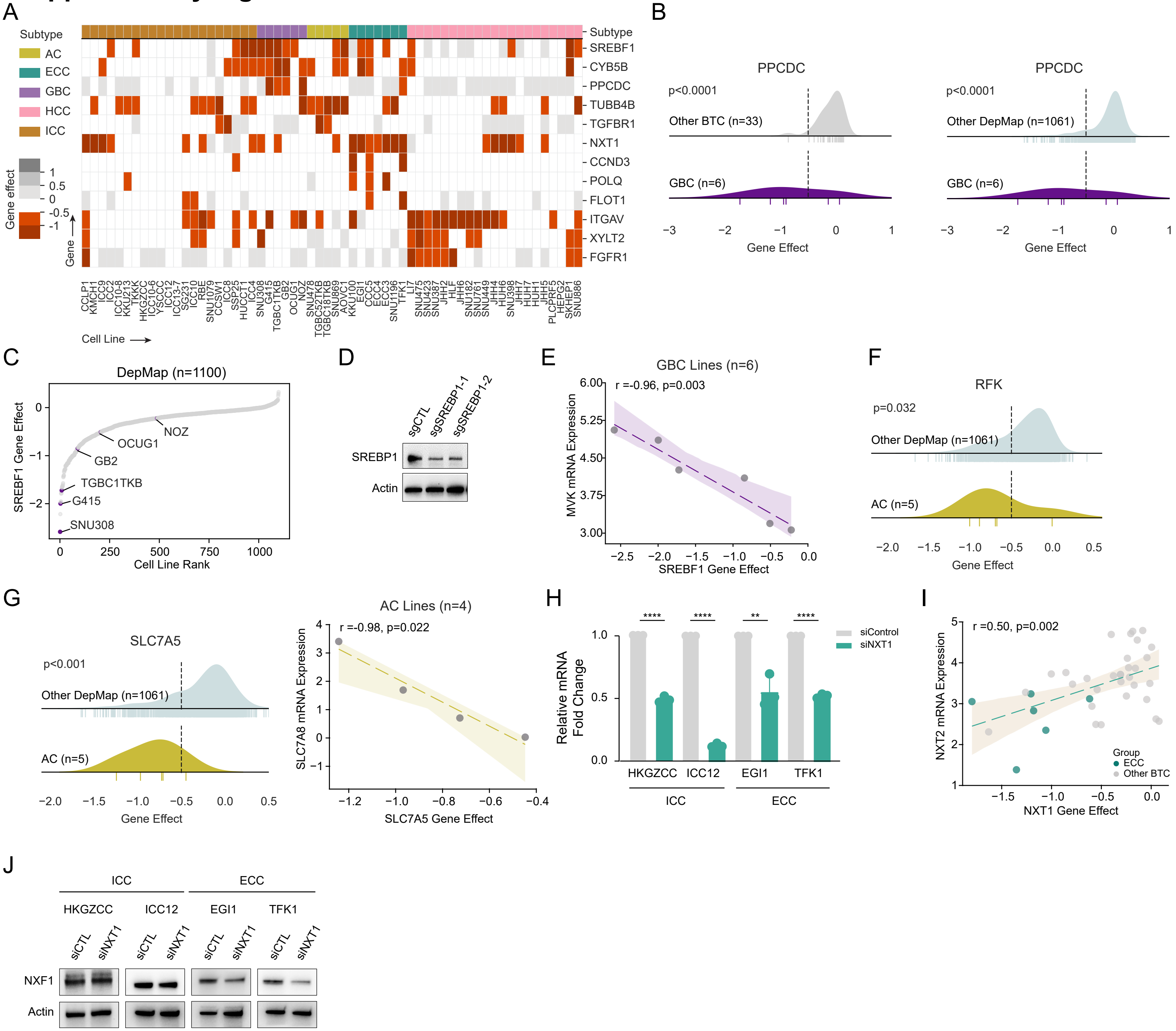

### Supplementary Figure5.pdf

#### Supplementary Figure 5

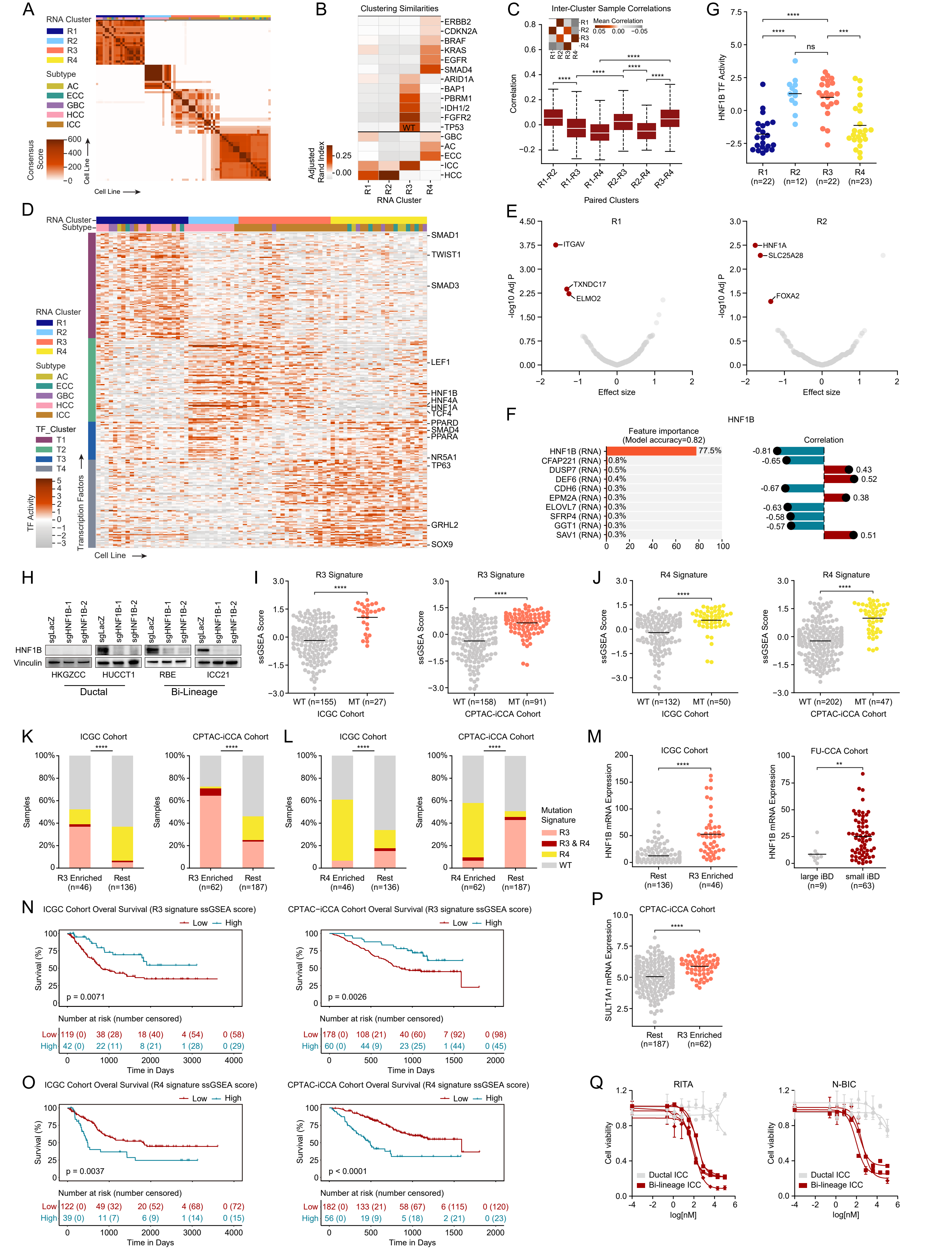

### Supplementary Figure6.pdf

#### Supplementary Figure 6

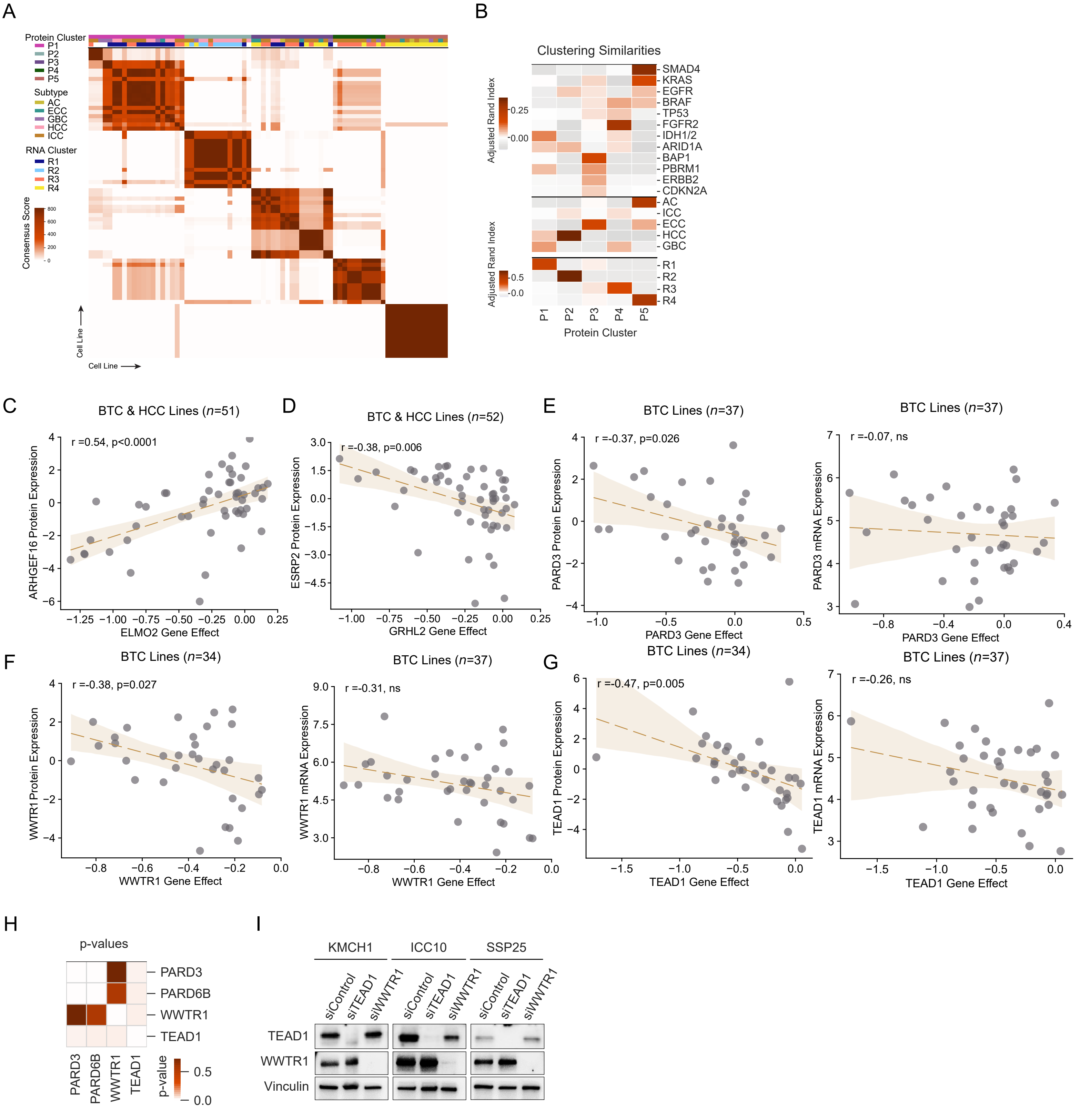

### Supplementary Figure7.pdf

#### Supplementary Figure 7

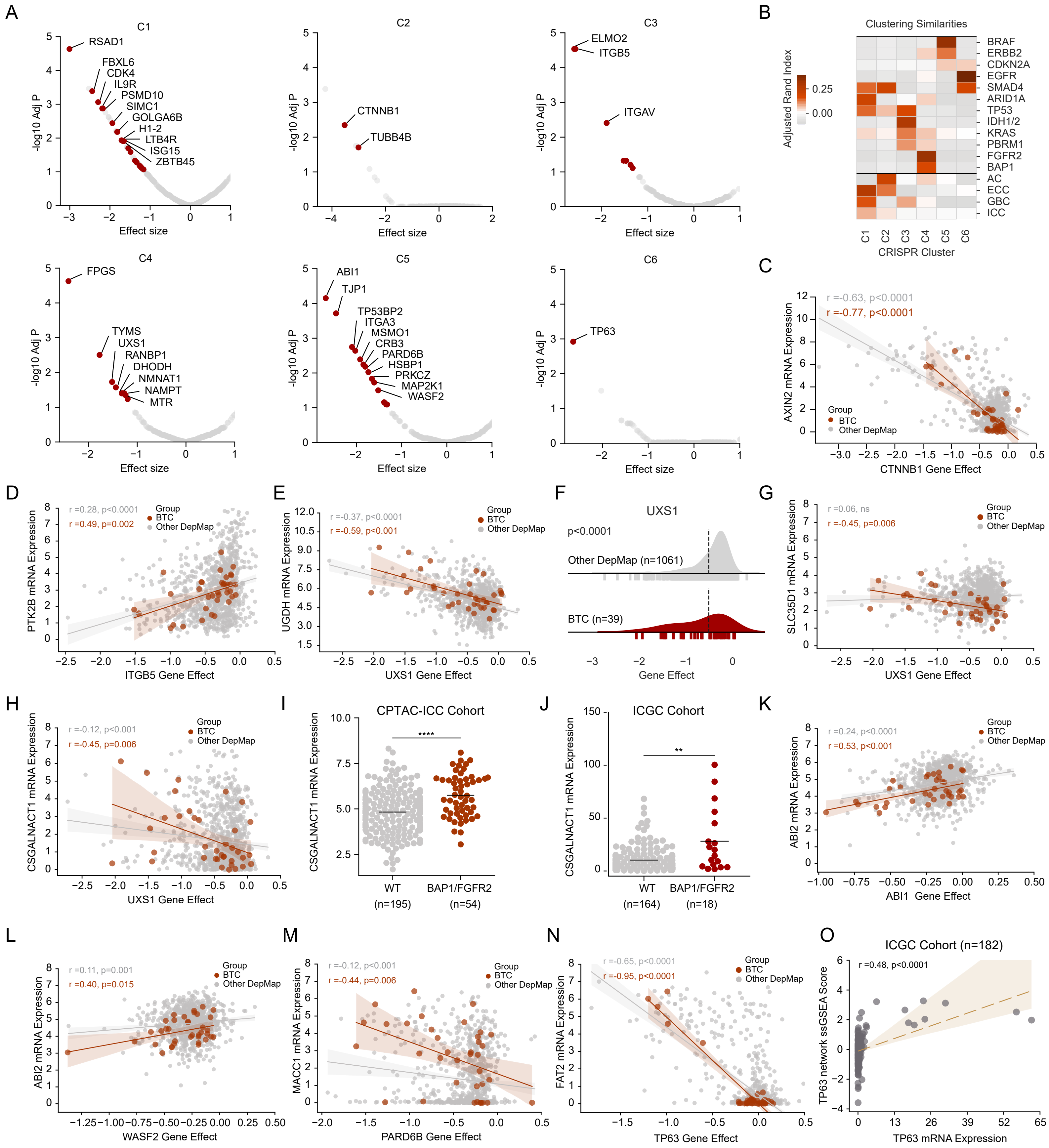
